# *RRM2B* is frequently amplified across multiple tumor types: non-oncogenic addiction and therapeutic opportunities

**DOI:** 10.1101/2020.09.10.291567

**Authors:** Waleed Iqbal, Elena V. Demidova, Samantha Serrao, Taha ValizadehAslani, Gail Rosen, Sanjeevani Arora

## Abstract

*RRM2B* plays a crucial role in DNA replication, repair and oxidative stress. While germline *RRM2B* mutations have been implicated in mitochondrial disorders, its relevance to cancer has not been established. Here, using TCGA data, we investigated *RRM2B* alterations in cancer. We found that *RRM2B* is highly amplified in multiple tumor types, particularly in *MYC*-amplified tumors, and is associated with increased *RRM2B* mRNA expression. We also observed that the chromosomal region 8q22.3–8q24, is amplified in multiple tumors, and includes *RRM2B*, *MYC* along with several other cancer-associated genes. An analysis of genes within this 8q-amplicon showed that cases that have both *RRM2B*-amplified along with *MYC* have a distinct pattern of amplification compared to unaltered cases or cases that have amplifications in *RRM2B* or *MYC* only. These other 8q-proteins were shown to interact functionally within the RRM2B network of DNA repair, hypoxia and apoptosis regulating proteins. Notably, *RRM2B*-amplified tumors are characterized by mutation signatures of defective DNA repair and oxidative stress, and in some cancers also associated with poor clinical outcome. These findings suggest that some cancers may require RRM2B for cellular survival, providing novel therapeutic opportunities in these cancers.

## Introduction

RRM2B plays an important role in regulating replication stress, DNA damage, and genomic stability [1,2]. *RRM2B* encodes a small subunit of p53-inducible ribonucleotide reductase (RNR). The RNR is a heterotetrametric enzyme responsible for the *de novo* conversion of ribonucleotide diphosphates into the corresponding deoxyribonucleotide diphosphates for DNA synthesis, thus playing an important role in maintaining deoxyribonucleotide pools [3]. The large subunit of the RNR complex consists of a dimer of the RRM1 protein, while the small subunit dimer is either RRM2 or RRM2B (varies depending on cellular conditions). P53-dependent induction of RRM2B expression by hypoxia leads to the exchange of the small RNR subunit from RRM2 to RRM2B, forming a new RNR complex that drives basal DNA replication, reduces replication stress, and maintains genomic stability [4,5]. These known functions of RRM2B suggest that *RRM2B* alterations may play a role in tumorigenesis [1,2].

RRM2B is located on chromosome 8q (8q23.1 [6]; in 2018, annotation changed to 8q22.3 (https://useast.ensembl.org/)). Germline missense and loss of function mutations in *RRM2B* have been associated with mitochondrial depletion syndrome (MDS), with distinct but variable clinical phenotypes [7,8]. At present, there are no known *RRM2B* germline alterations associated with cancer risk. However, somatic changes in *RRM2B*, including most typically amplifications, have been observed in multiple cancers including breast, liver, lung and skin cancer [9]. In a survey of the COSMIC database, *RRM2B* emerged as the most highly amplified DNA repair gene [9]. Additionally, TCGA studies of ovarian, breast, liver, and prostate cancer, have found that cases with *RRM2B* copy number variations (CNV) (amplifications and deletions) have decreased overall survival (OS) [9]. Similarly, increased metastasis and poor prognosis were correlated with *RRM2B* overexpression in head and neck cancer [10], esophageal cancer [11] and lung sarcomatoid carcinoma [12]. Another study noted that elevated expression of *RRM2B* is associated with better survival in advanced colorectal cancer [13].

*RRM2B* amplification may be driven by selection of *RRM2B* function, or *RRM2B* may be amplified as a passenger, concurrent with selection for a nearby gene with a driver activity in cancer. In breast cancer, multiple genes localized in the 8q12.1 - 8q24.23 interval were found to be amplified, including *RRM2B* [14]. Most *RRM2B* amplifications are accompanied by *MYC* amplifications, and these two genes are located in close proximity [15]. However, *RRM2B* amplifications also occur independent of *MYC* amplifications, albeit at a lower frequency [16]. While multiple studies have observed the amplification of the 8q region, currently the frequency and specificity of these amplifications is not known, and more specifically, the consequence of *RRM2B* amplification with *MYC* or as independent from *MYC* amplification is not clearly understood [14,17–26].

Here, using TCGA data, we found that *RRM2B*-amplified tumors not only exhibit increased *RRM2B* expression in multiple cancers (such as breast, ovarian, and head and neck cancer), but also exhibit distinct mutation signatures. Comparison of tumors bearing single gene amplifications of *RRM2B*, *MYC* versus co-amplifications (*RRM2B* and *MYC*) indicated that *RRM2B* amplifications may independently impact clinical outcomes in some cancers. An analysis of tumors with *RRM2B* and *MYC* co-amplifications or *RRM2B* alone showed that several genes in the 8q22 – 8q24 region are highly amplified. Additionally, RRM2B and these 8q-proteins interact within the same cell regulatory mechanisms of DNA damage response and repair, hypoxia and apoptosis. Based on these results, we hypothesize that while *MYC* may be the cancer driver, co-amplification of *RRM2B* and other 8q-genes may be relevant for cancer cell survival. These finding present opportunities for novel therapeutic targeting strategies (such as those targeting DNA damage response and repair) for tumors carrying *RRM2B* alterations.

## Materials and Methods

### Analysis of alteration frequencies using TCGA studies

TCGA tumor data was accessed using the cBioPortal website (http://www.cbioportal.org) [60]. Tumors that have been profiled for mutations as well as copy number variants from 30 TCGA studies were downloaded on 6/21/2018. For cancer types with the highest frequency of *RRM2B* amplifications, cases were segregated based on co-occurrence of *TP53* mutations or *MYC* amplifications. The *TP53* alterations were all mutations (missense, and truncating). The truncating mutations in TCGA are frameshift deletions, frameshift insertions, nonsense, splice site. All mutations were analyzed for co-occurrence, including those that were homozygous or heterozygous.

### RNA expression analysis

RNA seq. V2 RSEM data was downloaded from OV, BRCA and HNSC studies on 11/7/2018, and 3/19/2020. The RNA seq. data was only analyzed for tumors that were also profiled for copy number variants. Cases were separated based on copy number variants type: deep deletion, shallow deletion, diploid, gain and amplification. For *RRM2B* there was only a single case of deep deletion out of all (n= 2181 cases from all studies), observed in BRCA, and was removed. For *CCNE2, EI3FE, MTDH, MYC, RAD21, TP53INP1* and *YWHAZ* - there were no cases of deep deletion for any of the genes in the HNSC study, but there was one case each in *EI3FE, RAD21* and *YWHAZ* with deep deletions in the BRCA study, and also one case each in *EI3FE* and *MTDH* as well as two cases in *RAD21* for the OV study. In the supplementary figure 2, the box plot represents log2 mRNA expression values from RNA seq. V2 RSEM from cases with different copy number variants categories and statistical significance was tested in Graphpad Prism 8.0 using Mann-Whitney Non-Parametric T-test [62,63]. Additionally, plots of log2 mRNA expression values based on relative linear copy number values were used to test for correlation between increasing copy number change and mRNA expression, then Pearson coefficient [64] and log rank [65] p-values for each cancer type were calculated. Graphpad Prism 8.0 was used for all statistical analysis. For *CCNE2, EI3FE, MTDH, MYC, RAD21, TP53INP1 and YWHAZ*: the data were analyzed as above, but are presented as supplemental tables with expression data and statistics.

### Mutation signature analysis

Tumor whole-exome sequence data for TCGA Pan-Cancer Atlas studies (for OV, BRCA, HNSC and LUAD), which includes spectra of individual tumors, were downloaded from the Synapse platform (synapse.org) on 11/5/2018. Next, IDs of subjects with different cancer types were downloaded from the cBioPortal website (cBioPortal.org), and subjects with amplification in *RRM2B* and *MYC* genes were selected. Presence or absence of these amplifications classified the subjects into four categories: 1) cases with amplification in both *RRM2B* and *MYC*, 2) cases with amplification in *RRM2B*, but not in *MYC*, 3) cases with amplification in *MYC*, but not in *RRM2B*, 4) and cases without amplification in both genes. For each specific cancer type, mutational spectra of each category were calculated by finding the average number of amplifications in each of the 96 mutation classes. To compare differences in these 96 mutation types, one-way ANOVA [66], two-way ANOVA [67,68], and Wilcoxon rank-sum test [67] were used to test for significant differences. In order to eliminate the possibility of false discoveries caused by multiple comparison, in each test, Benjamini–Hochberg correction [69] was applied to each group of 96 p-values (corresponding to 96 mutations). For each cancer type the patients were divided into the following mutation signature groups: 1) cases with amplification in both *RRM2B* and *MYC*, 2) cases with amplification in *RRM2B*, but not in *MYC*, 3) cases with amplification in *MYC*, but not in *RRM2B*, 4) and cases without amplification in both genes, and anova1, anova2 and Wilcoxon rank-sum functions in MATLAB R2018a were used to test for significance. Each signature type was determined to be significant only if its P-value was below 0.05 in the one-way ANOVA test. The two-way ANOVA was used to distinguish the impact of factors such as co-amplifications with *MYC*. Additionally, the Wilcoxon rank-sum test was also performed on the data as another analysis of significance.

### Survival analysis

Clinical data was downloaded for the OV, HNSC, LUAD and BRCA TCGA studies. Patients were divided into different groups based on having amplifications in *RRM2B* or *MYC*, in both or neither. The OV cases were all Stage 3 or 2 for *RRM2B* and *MYC*. The clinical data from each group was used to generate overall survival and disease/progression-free survival Kaplan-Meier curves [70] using GraphPad Prism 8.0, and each curve was compared with the other respective curves using the log-rank test [65] and significance was shown based on * for P-value <0.05, ** for P-value <0.01 and *** for P-value <0.001. The significance level was set at p<0.05. Additionally, Wilcoxon tests were also used [71], and significance was shown based on * for P-value <0.05, ** for P-value <0.01 and *** for P-value <0.001. The significance level was set at p<0.05.

### Analysis of amplifications in 8q-genes

Data previously used for TCGA alteration analysis (cbioportal.org) were downloaded on 6/22/18 for our analysis of amplifications in 8q-genes. TCGA studies were used for BRCA, OV and HNSC cancers. The genes used for analysis are located on chromosome 8 from the region 8q11.2 – 8q24.3 and have been associated with cancer according to the Cancer Index (www.cancerindex.org). The gene names, their chromosomal location, and nucleotide start positions are in **Supplementary Table 1**. The amplification pattern of these other 8q-genes was compared in cases with co-amplifications of *MYC* and *RRM2B*, cases with single amplifications or unaltered cases. A univariate analysis was conducted to calculate the positive or negative linear trend for the frequency of gene amplification, assessed at 95% confidence interval. The Pearson correlation coefficient and p-values were computed by using PROC CORR in SAS 9.4. The significance level was set at p<0.05.

### Interaction and pathway enrichment analysis

Data for the RRM2B interaction network was downloaded from the BioGRID database [72] (10/25/2018) and analyzed by Cytoscape v. 3.6.1 [73]. Several interactions were added manually based on literature findings (RRM2B-FOXO3 [34], RRM2B-P21 [32], RRM2B-TP73 [74], RRM2B-E2F1 [33], RRM2B-MEK2 [75]).

WebGestalt (WEB-based GEne SeT AnaLysis Toolkit) was used for gene set enrichment analysis to extract biological insights from the genes of interest [40]. The online WebGestalt tool was used, and an over representation analysis was performed. The enrichment results were prioritized by significant p-values, and FDR thresholds at 0.01.

## Results

### RRM2B is highly amplified in multiple cancers, with an amplification frequency similar to MYC, while alterations in other RNR genes are infrequent

Using cancer cases from TCGA, we observed that, *RRM2B* is frequently amplified, with the highest percentage observed in ovarian, breast, bladder, and liver cancers (21.54% - 14.5%), and a lower rate of amplifications in multiple other cancers (14% to 0.6%) (Figure 1A). Deletions of *RRM2B* were only observed in Non-Hodgkin’s lymphoma (~2%). In addition to amplifications, a low frequency of mutations (<2%) were observed in lung, endometrial, esophagogastric, cervical and pancreatic cancers (**Figure 1A**). Mapping of somatic *RRM2B* mutations to the RRM2B protein structure indicated the mutations are present in multiple regions of the protein and do not always cluster at defined functional domains/regions [27–29], unlike the mutations observed in mitochondrial disorders, which typically result in reduced or eliminated function of the RRM2B protein [7,8] (**Figure 1B**).

**Figure 1.**
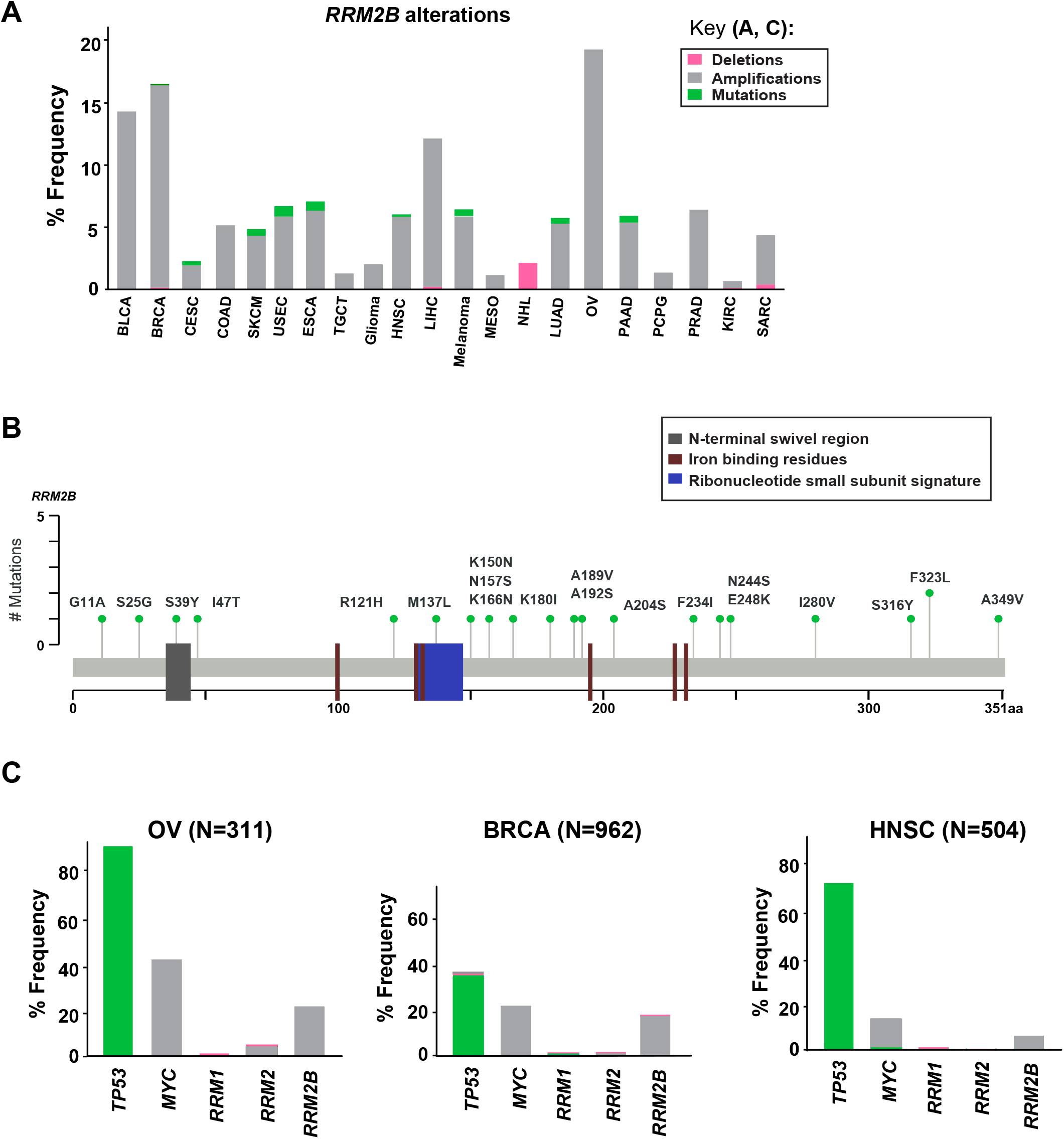
Somatic alterations in *RRM2B* from cbioportal.org. **A**: **Somatic alterations in *RRM2B* across TCGA studies:** BLCA (Bladder Urothelial Carcinoma), BRCA (Breast invasive carcinoma), CESC (Cervical squamous cell carcinoma and endocervical adenocarcinoma), COAD (Colon adenocarcinoma), SKCM (Skin Cutaneous Melanoma), USEC (Uterine Corpus Endometrial Carcinoma), ESCA (Esophageal carcinoma), TGCT (Testicular Germ Cell Tumors), HNSC (Head and Neck squamous cell carcinoma), LIHC (Liver hepatocellular carcinoma), MESO (Mesothelioma), NHL (Non-Hodgkin lymphoma), LUAD (Lung adenocarcinoma), OV (Ovarian serous cystadenocarcinoma), PAAD (Pancreatic adenocarcinoma), PCPG (Pheochromocytoma and Paraganglioma), PRAD (Prostate adenocarcinoma), KIRC (Kidney renal clear cell carcinoma), SARC (Sarcoma). Amplifications (grey), mutations (green) and deletions (pink) are represented as percent frequency. **B**: ***RRM2B* mutations in TCGA**. 2D RRM2B protein stick figure showing the important domains of RRM2B (N-terminal swivel region, required for dimer stability, grey; ribonucleotide small subunit signature [conserved region in catalytic site between RRM2 and RRM2B], blue, and iron-binding residues required for catalytic activity, red) and the number of somatic mutations in RRM2B. **C**: **Frequency of *TP53, MYC, RRM1, RRM2* and *RRM2B* alterations** in ovarian (OV), breast (BRCA) and head and neck (HNSC) cases.

Since RRM2B is regulated by TP53 and is typically co-amplified with *MYC* (**Figure 1**), we next compared the frequency of *RRM2B* alterations with alterations in *TP53*, and *MYC*. We also compared the frequency of *RRM2B* alterations along with the members of the RNR complex (*RRM1, RRM2* [30]) (**Figure 1C**). As expected, *TP53* was the most altered gene in the studied cancers, however, *RRM2B* amplifications did not always significantly co-occur with *TP53*. Interestingly, most cases with *TP53* mutations (which were missense, and truncating mutations) did not have *RRM2B* amplifications (**Supplementary Figure 1A**). In comparison, a greater number of cases with MYC amplifications were observed to also have *RRM2B* coamplifications. For ovarian (OV) and head and neck squamous cell carcinoma (HNSC), this was half of all cases with *MYC* amplifications, while in breast cancer (BRCA) this was ~90% cases (**Supplementary Figure 1B**). Finally, *RRM1* and *RRM2*, the functional partners of RRM2B, were infrequently altered and did not co-occur with *RRM2B* amplifications (**Figure 1C**).

### Tumors with RRM2B amplifications exhibit increased RRM2B expression

Previous studies have not shown whether tumors carrying amplified *RRM2B* have increased RRM2B expression, which might directly impact its role in cancer. Multiple studies have observed that gene amplifications do not always lead to increased expression [31]. Thus, to confirm increased expression, we used mRNA expression data from three TCGA studies (OV, BRCA, HNSC), and found that tumors with gains and amplifications had significantly increased *RRM2B* mRNA expression in all three cancer types studied (**Supplementary Figure 2**). *RRM2B* amplifications also significantly correlated with an increase in *RRM2B* mRNA expression in ovarian and breast cancer, and with an increased trend in HNSC (OV: Pearson correlation=0.64, Log rank p-value=0.05; BRCA: Pearson correlation=0.53, Log rank p-value=0.04; HNSC: Pearson correlation=0.32, Log rank p-value=0.08) (**Supplementary Figure 2**).

### RRM2B amplification by itself or co-amplification with MYC is in an 8q-amplicon that is present in multiple cancer types

In multiple cancers, we identified that *RRM2B* and/or *MYC* were amplified as part of an amplicon with multiple other 8q-genes (**Supplementary table 1**, list of genes and gene ontology classification). We queried the amplifications in OV (**Figure 2**), BRCA (**Supplementary Figure 3**), and HNSC (**Supplementary Figure 4**) cancers in TCGA.

**Figure 2.**
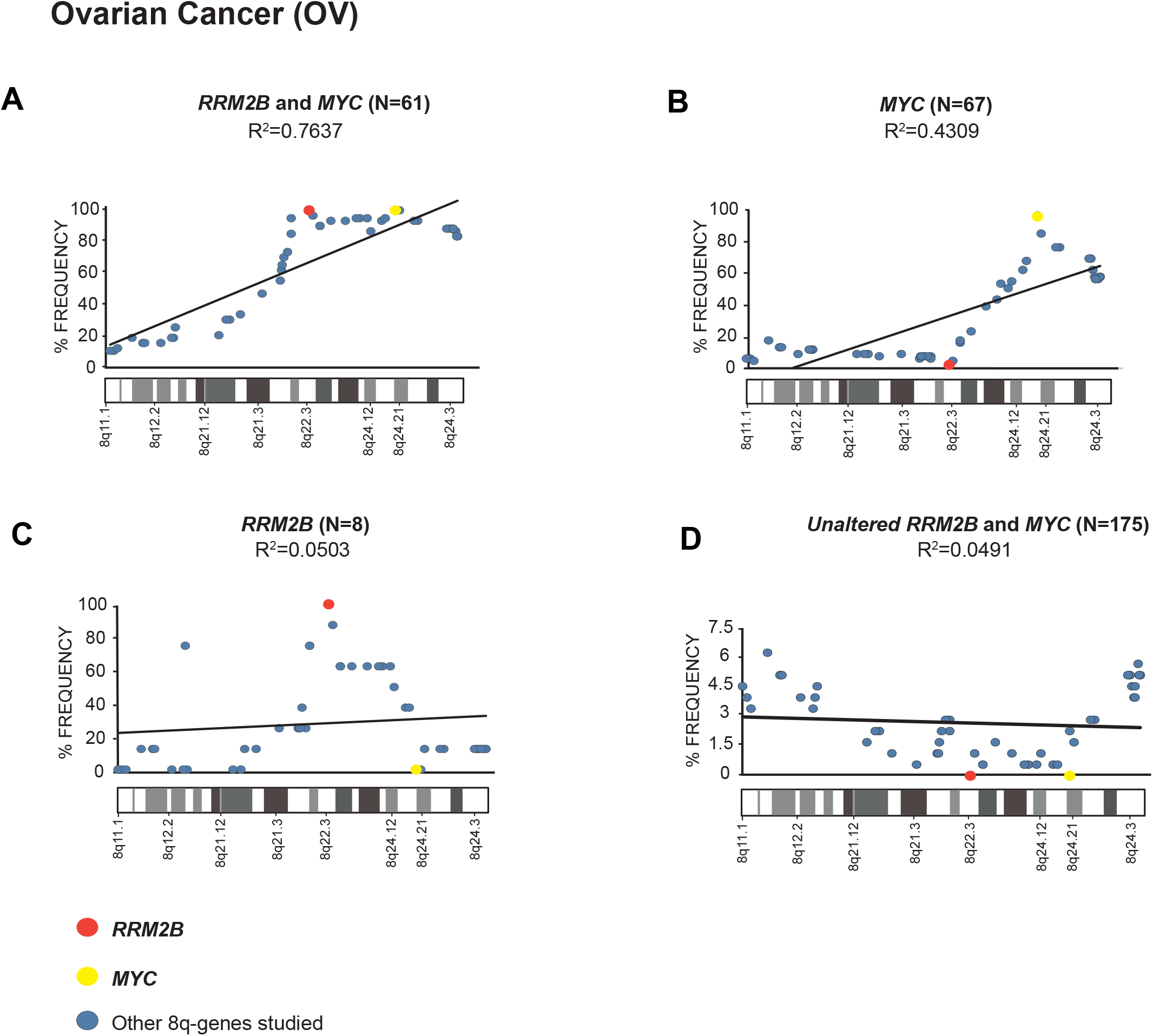
Amplification frequency of 8q-genes in ovarian cancer (OV). **A**: cases with coamplification of *RRM2B* and *MYC*; **B**: cases *MYC* only amplification; **C**: cases *RRM2B* only amplifications; **D**: cases with neither (unaltered) were plotted as percent of frequency for amplifications in various 8q-region genes relevant for cancer (see **Supplementary Table 1**). *RRM2B* (red circle), *MYC* (yellow circle), and other genes (blue circle). The Pearson correlation (R2 value) for the data points is represented by a black trend line.

We observed a strong Pearson correlation for amplifications in the 8q11-8q24 region in cases with *RRM2B* and *MYC* co-amplifications. The strongest correlation was observed for ovarian cancer (R^2^=0.7637, p-value <0.00001, **Figure 2A**). The cases with either *MYC* or *RRM2B* amplification alone (R^2^=0.4309, p-value= 0.000273, **Figure 2B**, and R^2^=0.0.0503, p-value= 0.905, **Figure 2C**) or those without either (R^2^=0.7637, p-value= 0.518, **Figure 2D**) showed weaker Pearson correlation. These amplifications observed (**Figure 2A-C**) are in the chromosome segment between *RRM2B* (8q22.3) and *MYC* (8q24.21). This amplicon contains 11 cancer relevant genes: *BALC, ANGPT1, EIF3E, EBAG9, TRSP1, RAD21, EXT1, TNFRSF11B, NOV, HAS2* and *RNF139* (**Supplementary table 1**).

Next, we examined the overall amplification frequency of 8q genes using TCGA data, without segregating cases based on *RRM2B* and/or *MYC* amplifications. We found, that the 8q11.3-8q24 amplicon is present in multiple tumor types (**Figure 3**). A positive correlation was observed for increased amplifications in the 8q11-8q24 region, which includes *RRM2B* and *MYC* (Pearson correlation range: R^2^ =0.8976-0.0568). The strongest correlations were observed for skin (SKCM, R^2^=0.8976, p-value <0.00001), pancreatic adenocarcinoma (PAAD, R^2^=0.8714, p-value <0.00001), ovarian cancer (OV, R^2^=0.8582, p-value <0.00001), liver hepatocellular carcinoma (LIHC, R^2^=0.7814, p-value <0.00001), and esophageal cancer (ESCA, R^2^=0.7376, p-value <0.00001) (**Figure 3**). All other correlations were significant at p-value <0.00001, except for BLCA (p=0.525). Finally, we also observed that mRNA expression of multiple genes in the amplicon was also significantly increased (**Supplementary Table 2**).

**Figure 3.**
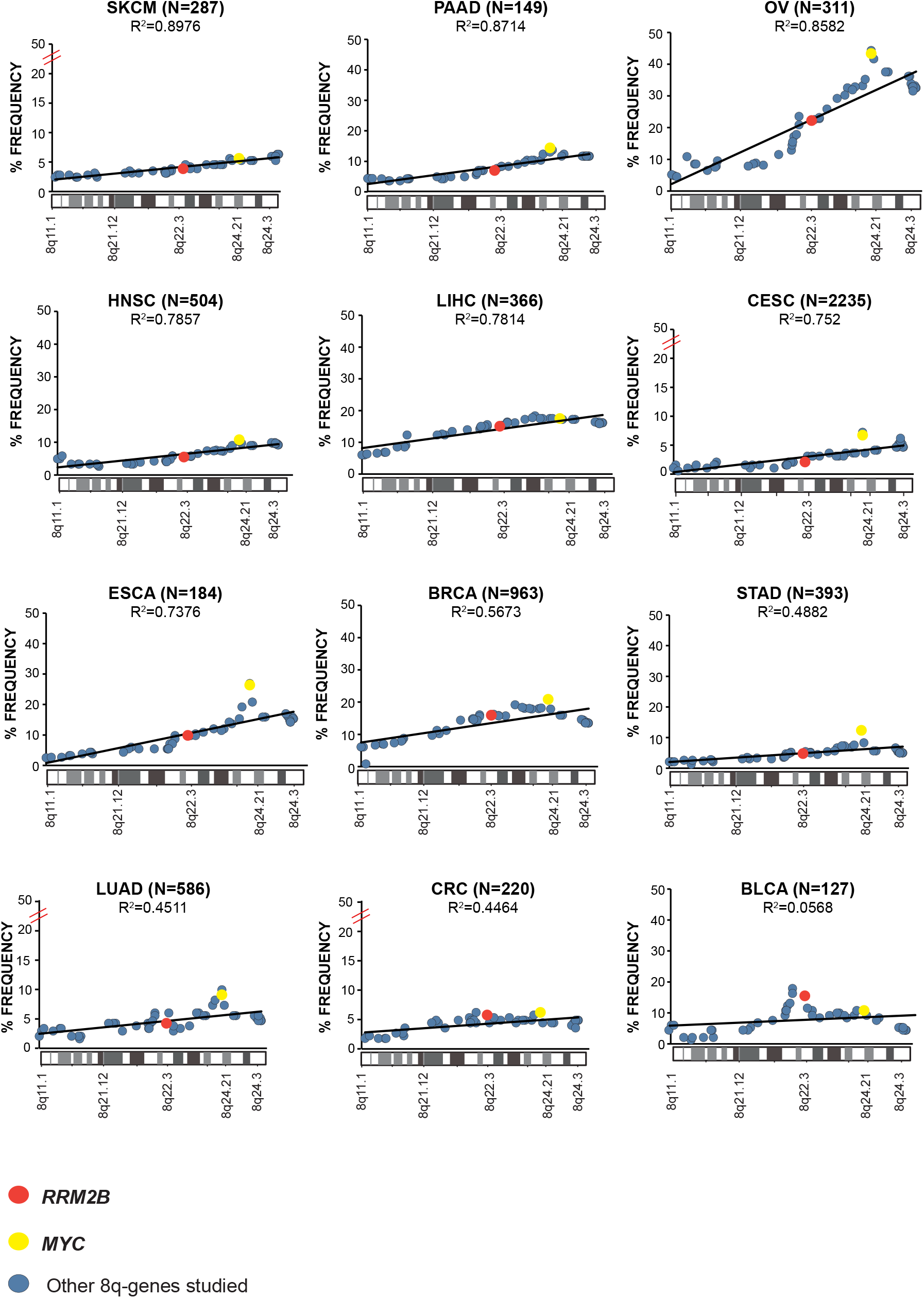
Amplification frequency of cancer relevant 8q-genes in Skin Cutaneous Melanoma (SKCM), Pancreatic adenocarcinoma (PAAD), Ovarian serous cystadenocarcinoma (OV), Head and Neck squamous cell carcinoma (HNSC), Liver hepatocellular carcinoma (LIHC), Cervical and endocervical cancers (CESC), Esophageal carcinoma (ESCA), Breast invasive carcinoma (BRCA), Stomach adenocarcinoma (STAD), Lung adenocarcinoma (LUAD), Colon adenocarcinoma (COAD), Bladder urothelial carcinoma (BLCA) amplifications were plotted as percent of frequency for amplifications. *RRM2B* (red circle), *MYC* (yellow circle), and other genes (blue circle). The Pearson correlation (R^2^ value) for the data points is represented by a black trend line.

### RRM2B protein interaction network includes co-amplified 8q-amplicon genes

Since the expression of *RRM2B, MYC* and several other 8q-amplicon genes was increased, we next tested if the products of these genes within the 8q-amplicon interact with RRM2B. RRM2B is regulated by p53 and p21 [32] and transcriptional factors, such as, E2F1[33]. E2F1 regulates RRM2B in the absence of p53, and FOXO3 [34]. It binds the RRM2B promoter and induces its expression. Amplification in RRM2B may further impact the interacting proteins, which include proteins such as ATM, ATR, CHEK1, CHEK2, MDM2. These proteins are important for cell regulatory mechanisms such as DNA damage and response (DDR) pathway, cell cycle and apoptosis. Hypoxia pathway participants, VHL and HIF1α, interact with the above-mentioned DDR proteins, and with p53 and Myc.

We next found that several 8q22-24 gene products (**Figure 4**, in light purple) also interact or intersect with the proteins in the RRM2B network. For example, YWHAZ or 14-3-3 protein zeta/delta, is an adapter protein, which binds with phosphorylated FOXO3, which retains phosho-FOXO3 in the cytoplasm and prevents its activity as a transcription factor, and potentially prevents apoptosis [35]. Under certain conditions, such as irradiation, ATM-dependent activation of p53 involves dephosphorylation of p53 Ser376 residue, leading to formation of a binding site for YWNAZ, thus, increasing the affinity of p53 to bind with regulatory parts of genes such as *CDKN1A, GADD45, MDM2*, providing transcriptional regulation [36]. Another 8q-gene product found in the RRM2B network is EIF3E, which is required for several steps in the protein synthesis initiation. IF3E binds with the 40s ribosomal subunit to facilitate recruitment of proteins to initiate protein, and these complexes participate in ribosomal disassembly [37]. EIF3E interactions with ATM and BRCA1 are essential for execution of DDR, and altered EIF3E has been observed in breast cancer [38]. EIF3E has been suggested to play a role in tumor promotion (due to its interactions with MYC, an oncogene) or tumor suppression (due to interactions with NDRG1, a tumor suppressor protein and a target of MYC) [39].

**Figure 4.**
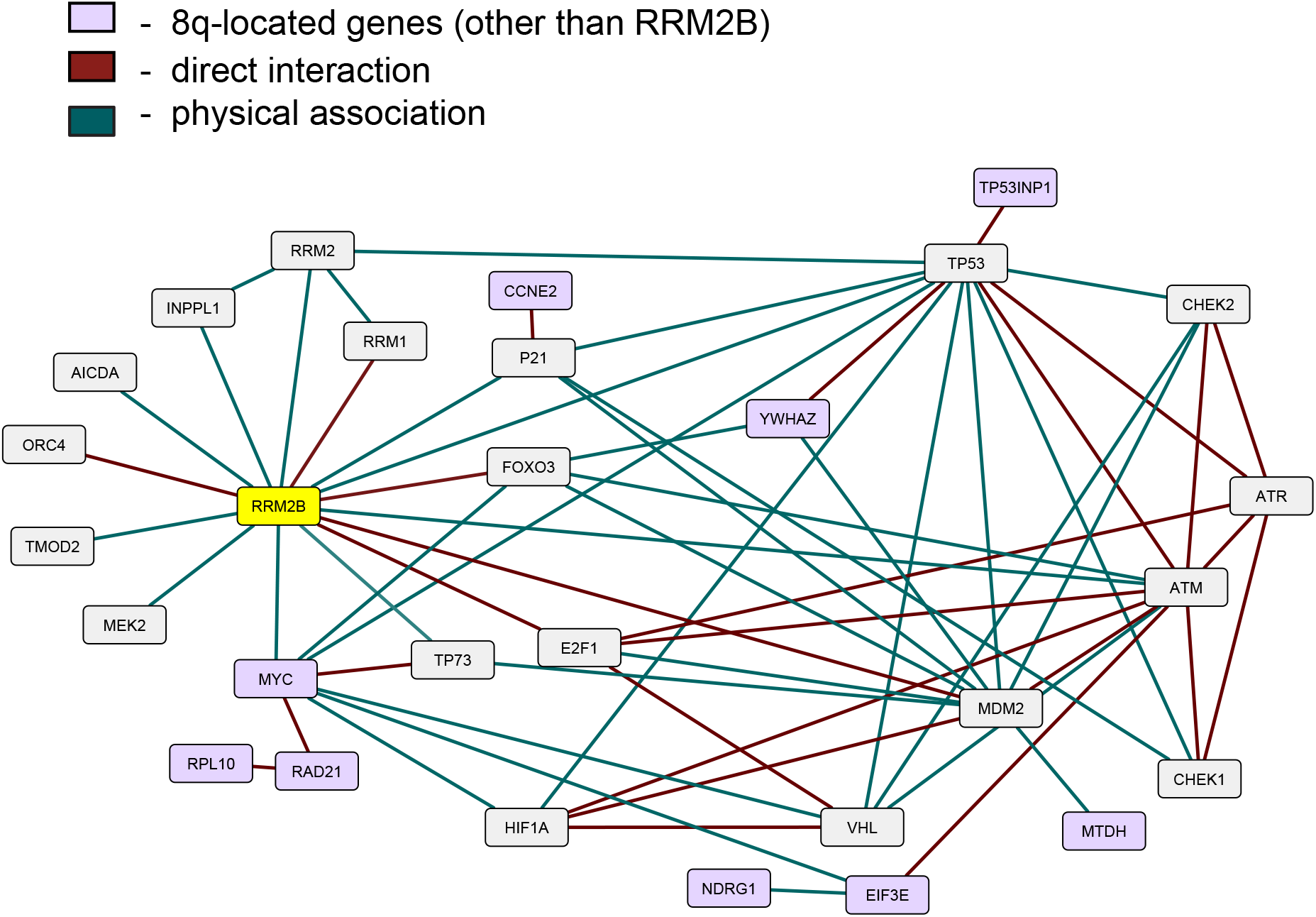
An interactive network of *RRM2B* and related genes. The network represents proteins that functionally interact or intersect with RRM2B (yellow) or other 8q-gene products (purple). Red lines represent direct interaction, and green lines – physical association (interactions classified according to the BioGRID database annotation).

Finally, we performed a pathway enrichment analysis for all the interacting genes presented in **Figure 4**. Using WebGestalt tool [40], we found that the most significantly enriched pathways were signal transduction mediated by p53, response to DNA damage and other environmental stimuli, cell cycle checkpoints, DNA replication, response to oxygen levels and apoptotic signaling. The results are presented in **Supplementary Table 3**. Overall, these data suggest that amplifications in the 8q-region, observed in multiple cancers, may impact cancer cell survival due to their involvement and intersection in important cellular pathways.

### *RRM2B amplifications correlate with clinical outcome in* ER+PR+HER2+ *breast cancer*

Previous studies have suggested that *RRM2B* alterations may impact patient prognosis in ovarian, breast, HNSC, and lung cancers [9–12]. However, these studies did not segregate cases based on relevant alterations, such as *MYC*. Here we compared survival differences in cases segregated based on amplifications in *RRM2B* alone, *RRM2B* with *MYC* or *MYC* alone in BRCA, OV, HNSC, and lung cancer (**Figure 5**, and **Supplementary Figure 5**).

**Figure 5.**
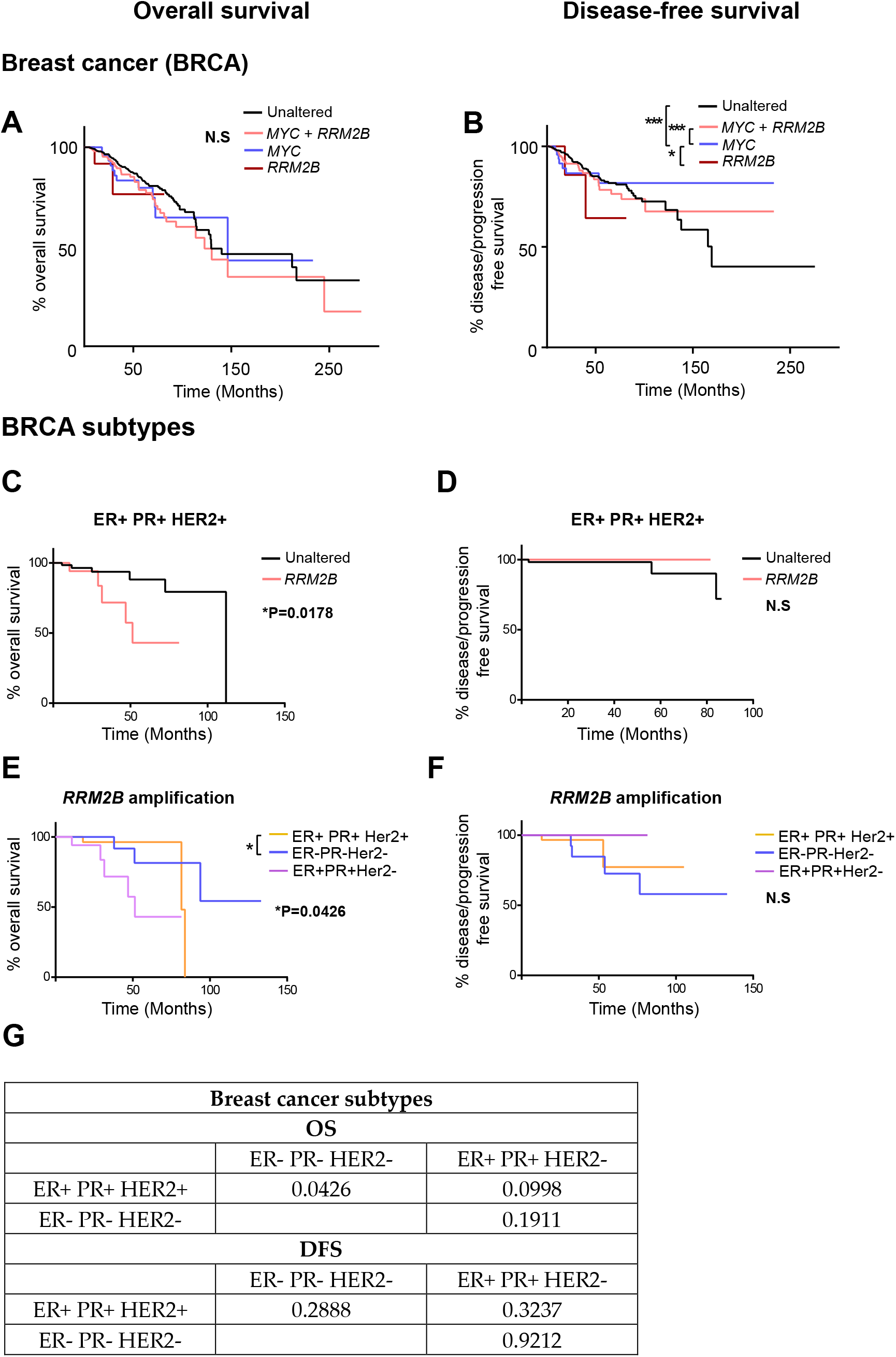
Kaplan-Meier curves for overall survival (OS) and disease/progression-free survival (DFS) in breast cancer (BRCA) (**A, B**) and its subtypes (**C-G**) studies with *RRM2B* amplifications and/or *MYC* amplifications. **A**: **OS in BRCA cases**. The cases that were unaltered for both genes (black, n=737), cases with *RRM2B* amplifications (red, n=14), cases with *MYC* amplifications (blue, n=62) and co-amplifications (pink, n =210) were plotted. **B**: **DFS in BRCA cases**. The cases that were unaltered for both genes (black, n=678), cases with *RRM2B* amplifications (red, n=12), cases with *MYC* amplifications (blue, n=55) and coamplifications (pink, n=184) were plotted. **C: OS in ER+ PR+ HER2+ BRCA cases**. The cases that were unaltered for RRM2B (black, n=68), cases with *RRM2B* amplifications (salmon, n=18). **D: DFS in ER+ PR+ HER2+ cases**. The cases that were unaltered for *RRM2B* (black, n=63), cases with *RRM2B* amplifications (red, n = 13). **E: OS in BRCA subtypes with *RRM2B* amplifications**. ER+ PR+ HER2+ cases with amplifications in *RRM2B* (purple, n=18), ER- PR- HER2- cases with amplifications in *RRM2B* (blue, n=26), and ER+ PR+ HER2- cases with amplifications in *RRM2B* (peach, n=36). **F: DFS in BRCA subtypes with *RRM2B* amplifications**. ER+ PR+ HER2+ cases with amplifications in *RRM2B* (purple, n=13), ER- PR- HER2- cases with amplifications in *RRM2B* (blue, n=25), and ER+ PR+ HER2- cases with amplifications in *RRM2B* (peach, n=34). The plots were compared using Log-rank test and significance is shown as follows: P-value *<0.05, ** for P-value <0.01 and *** for P-value <0.001. **G: P-values for the graphs E and F**.

In breast cancer, no significant differences in OS were observed (**Figure 5A**), which is a different result from a previous study in breast cancer [9]. However, in comparison to OS, significant differences were observed for DFS (**Figure 5B**). *MYC* amplification alone had the best DFS, compared to *MYC* and *RRM2B* co-amplified cases, and unaltered cases (p<0.0001). Additionally, *RRM2B* amplification cases tend to have worse DFS than *MYC* only cases (p=0.0519). To further explore the findings in breast cancer, we analyzed *RRM2B* amplifications separately in each of the three major subtypes of breast cancer (ER+PR+HER2+ (n=86), ER+PR+HER2- (n=334), and triple negative ER-PR-HER2- (n=100)). We observed that patients with ER+PR+HER2+ breast cancer and *RRM2B* amplifications had a significant decrease in OS (p=0.0178, **Figure 5C** and **5E**), with no difference in DFS (**Figure 5D, 5F**). Additionally, when comparing OS in all three breast cancer subtypes with *RRM2B* amplifications, ER+ PR+ HER2+ patients (n=18) had significantly worse OS than ER- PR- HER2- patients (n=26) (p=0.0426). The previous studies that observed worse OS in breast cancer, based on *RRM2B* amplification, did not consider subtype differences, and may have only included the major breast cancer subtype (ER+PR+HER2+).

In ovarian cancer, individuals with *MYC* amplification alone were observed to have a significantly better OS compared to those with *MYC* and *RRM2B* co-amplification (p=0.0019), or those that were unaltered for both genes (p=0.014) (**Supplementary Figure 5A**, see legend for case numbers). There were not enough *RRM2B* cases alone to evaluate statistically, but the data trended towards worse OS. No significant differences were observed in OV cancer for disease-free survival (DFS) (**Supplementary Figure 5B**). In lung cancer (non-small cell lung cancer or LUAD), significant difference was only observed in DFS for *RRM2B* only cases compared to unaltered cases. *RRM2B* amplifications alone had significantly worse DFS compared to unaltered cases (p=0.0057). Overall, there were not enough cases for statistical evaluation. For OS, there were only 7 cases with amplifications in both *RRM2B* and *MYC*, 15 cases with amplifications in *MYC* only, 6 cases with amplifications in *RRM2B* only, and 195 unaltered cases with no amplifications in either gene (**Supplementary Figure 5C**). For DFS, there were only 7 cases with amplifications in both *RRM2B* and *MYC*, 13 cases with amplifications in *MYC* only, 6 cases with amplifications in *RRM2B* only and 162 unaltered cases with no amplifications in either gene (**Supplementary Figure 5D**). Finally, no significant differences were observed based on *RRM2B* amplification status with or without *MYC* amplification, in OS and DFS in HNSC (**Supplementary Figure 5E** and **5F**). Additionally, p-values for all comparisons (log-rank test) in all the above cancer types is available in **Supplementary Table** 4.

### RRM2B amplifications exhibit a distinct tumor mutation signature but do not impact tumor mutation burden

As RRM2B plays an important role in DNA synthesis and maintaining genomic stability, we also tested whether *RRM2B* alterations impact tumor mutation burden (TMB). TMB is a biomarker of defective cellular pathways and mutagenic processes in the cell, and is emerging as a potential biomarker of cancer therapies, such as immune checkpoint blockade [41]. We first compared TMB (in OV, BRCA, and HNSC) in cases with and without *RRM2B* amplification, and did not find any differences in mutation rates, with similar rates observed for both (**Supplementary Table 5**).

Recently, distinct mutation signatures have been associated with defects in certain genes or pathways, and with certain endogenous and exogenous exposures [42]. Here, we tested whether *RRM2B* amplifications are associated with specific tumor mutational signatures. For this analysis, we used the recent PanCancer Atlas data, reporting mutational spectra for 9493 individual tumors, including 926 BRCA, 384 OV, and 461 HNSC and 524 LUAD [42]. Interestingly, breast cancer cases with *RRM2B* amplification alone were significantly associated with T>C and T>A mutations (P<0.05, **Figure 6 and Supplementary Table 6**).

**Figure 6.**
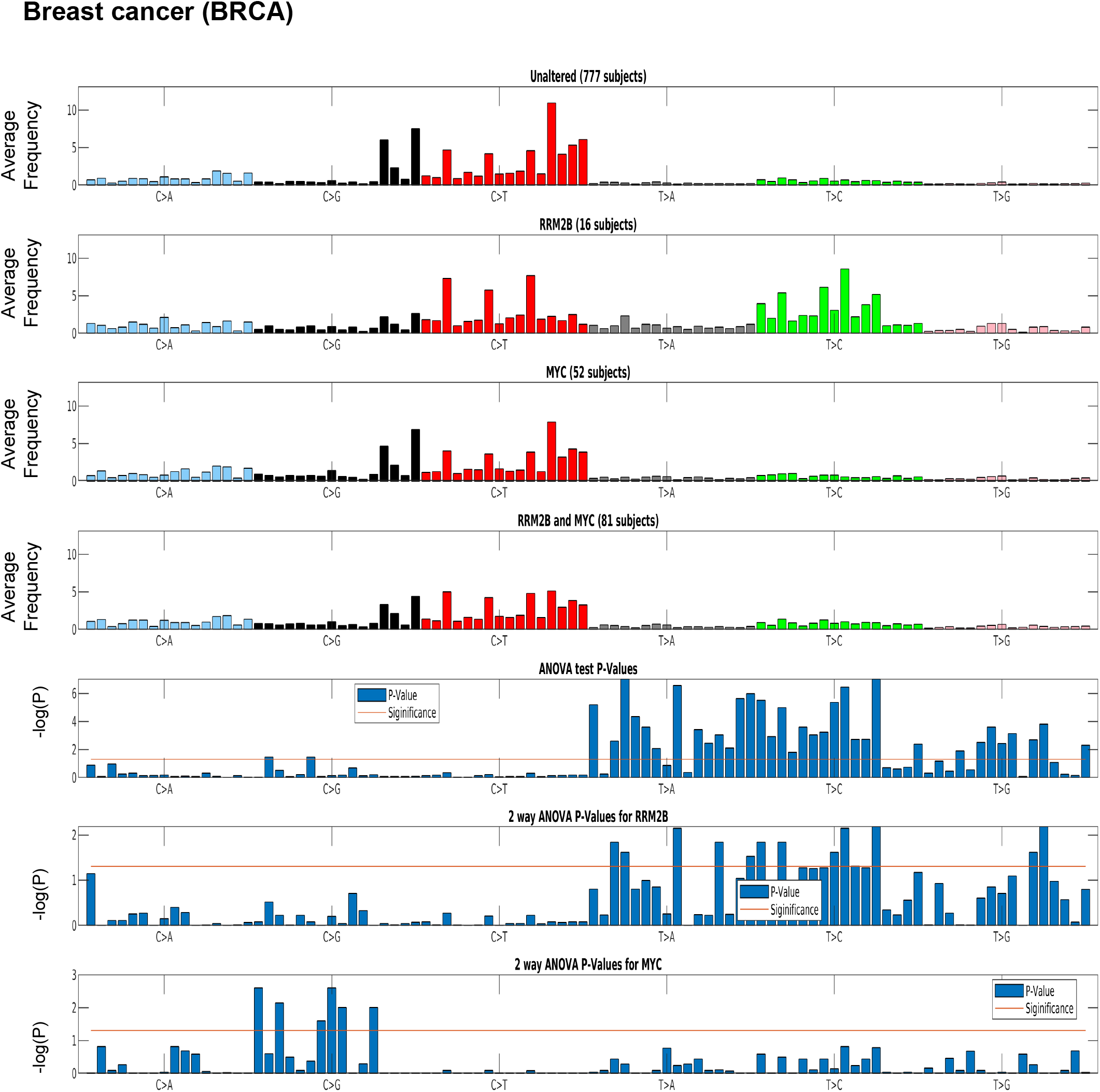
Mutation signature in breast cancer (BRCA) cases segregated by amplification type. One-way ANOVA (*RRM2B* amplifications only versus other groups) and two-way ANOVA (included group with *RRM2B* and *MYC* co-amplifications) analysis showed that the T>C and T>G mutations are statistically significant. **Top panels**: tumor whole-exome sequence data from the PanCancer Atlas studies was used to calculate the average frequency of the 96 trinucleotide context mutations in each group: unaltered cases, cases with only *RRM2B* amplifications, cases with only *MYC* amplifications, and cases with amplifications in both. **Bottom panels:** The statistical significance of each comparisons is represented by ANOVA tests as −log10 (P-value) for each of the 96 trinucleotide context mutations. The −log10 (P) visualizations are provided for: one-way ANOVA comparing *RRM2B* only group to the all other groups, a two-way ANOVA comparing all groups with *RRM2B* or *MYC* amplifications. −log10 (P) - the taller the bars are, the lower is the P-value and when the bars exceed the red line, P-values are less than 5%. P-values of each subfigure have been corrected using Benjamini–Hochberg procedure.

A similar signature was observed in HNSC, but not in OV (**Supplementary Figure 6**). The HNSC and ovarian cancer signature could not be tested for significance as only few cases were present in the PanCancer Atlas data. For LUAD, a distinct signature of C>A mutations (significant by one-way ANOVA and two-way ANOVA tests as above, p<0.05; **Supplementary Figure 7, and Supplementary Table 6**) was observed for cases with only *RRM2B* amplifications.

The observed mutation signatures for *RRM2B* amplifications were most similar to those that have been recently described for defects in distinct DNA replication and repair pathways [42]. The mutation signatures observed have been associated with defective DNA base excision repair, and DNA mismatch repair. We also observed mutation signatures for presence of reactive oxygen species that can lead to extensive DNA base damage [43].

## Discussion

Alterations in genes involved in DNA replication and/or repair pathways are known to increase genetic instability, and lead to cancer [9]. RRM2B is essential for DNA replication (nuclear and mitochondrial), repair, protection from oxidative damage, and overall maintenance of genetic stability. Despite these known functions, our current understanding of RRM2B is limited to its role in MDS. *RRM2B* alterations implicated in MDS are typically mutations that result in the reduction of mitochondrial dNTP pools [44]. The current understanding of *RRM2B* in cancer is very limited, and in this study, we provide a broad analysis of *RRM2B* alterations in cancer and their potential significance.

*RRM2B* is a major downstream target of p53, however, in contrast to p53 loss of function mutations or deletions, we observed that *RRM2B* is highly amplified across multiple tumor types. Interestingly, *RRM2B*, which is present on chromosome 8 is usually co-amplified with *MYC* oncogene (which is also on chromosome 8). Currently, from these observations it is challenging to establish *RRM2B* as a tumor promoter or suppressor, but these data may reflect potential addiction of cancer cells to *RRM2B*, and other genes in the 8q-amplicon observed in multiple cancers in this study. Our data warrant further investigation in cellular models to investigate role of *RRM2B* in cancer. In previous studies, RRM2B expression has been identified as a predictive marker of resistance to paclitaxel and gemcitabine in breast cancer, by using machine learning approaches [45]. Similarly, *RRM2B* has been identified as an essential metabolic gene for renal tumors, and suggested as a potential therapeutic target to disrupt cancer cell metabolism [46].

We also found, in multiple cancers, that tumors with co-amplified *RRM2B* and *MYC* exhibit significant amplification of 8q22.3 – 8q24 region of chromosome 8. Interestingly, a previous study in breast cancer found frequent *MYC* co-amplifications with multiple genes in the 8q chromosomal region. This study suggested that these co-amplifications may explain the aggressive phenotypes of these tumors [14]. In high-grade ovarian serous carcinomas, it was found that *MYC* was one of the most amplified genes, and patients with such copy number changes had better overall survival [47]. In this study it was not possible to address the problem of inter- and intra-tumor heterogeneity which has implications for therapeutic resistance [48,49]. However, supportively, a recent study in the highly heterogenous ovarian serous grade carcinomas consistently found amplifications (starting from 8q22.3 region) in ovarian tumor biopsies from different regions of the ovary, and also in the relapsed tumor tissue [50]. Future studies addressing inter- and intra-tumor heterogeneity in other cancer types will further highlight the potential clinical significance of amplifications in the 8q-region.

Amplification of *RRM2B*, along with other genes in the 8q-amplicon could reflect tumor non-oncogenic addiction in multiple tumors [51,52]. While *MYC* is the main driver for these tumors, the amplification *RRM2B* and the other 8q-genes may be relevant for cancer cell survival, with clinical and therapeutic implications. A previous study has observed prognostic significance of *RRM2B* alterations in nasopharyngeal carcinoma [12]. Our analysis of clinical outcomes revealed that based on the cancer type or subtype, co-alteration of *RRM2B* with *MYC*, or alone may significantly impact patient outcome in breast cancer. The clinical implications of these data will be further illuminated by large sample size, and analysis of other independent prognostic factors of overall survival. For example, absence of residual tumor postsurgery is a major prognostic factor in advanced ovarian cancer [53].

Finally, we observed that several genes within the 8q-amplicon also interact with the RRM2B-functional network. This suggests that amplifications in this region could have a profound effect on DNA replication, repair, cell cycle and hypoxia. Pathway enrichment analysis suggests the importance of 8q-region genes in response to oxygen levels, DNA replication, and DNA damage response. Future studies in cellular models are needed to validate the findings from the network analysis. Finally, at least in some cancers, *RRM2B* amplifications may either drive or participate in molecular events that lead to differential tumor mutation signatures. Here, the observed mutation signatures were relevant to the role of RRM2B in DNA replication, and repair pathways. A previous study in mice found that overexpression of RRM genes together with defective mismatch repair pathway can lead to increased mutagenesis, and carcinogenesis [54]. While a regulated expression of RRM2B reduces oxidative stress (in a p53-dependent manner) [55], the impact of *RRM2B* overexpression on the levels of reactive oxygen species is not known. The observation of mutation signatures associated with increased reactive oxygen species suggest that *RRM2B* overexpression may impede the oxidative stress pathway. The observed mutation signatures in *RRM2B*-amplified cases may assist in identifying cancer types or subtypes for precision-targeting of tumors. An example, is combining PARP inhibitors, with existing chemotherapies and irradiation [56–59].

Overall, this study provides an analysis of *RRM2B* alterations in multiple tumors, majorly reflected as *RRM2B* amplifications in conjunction with *MYC*, and other genes on chromosome 8. Future genome-wide studies of other cancer datasets are warranted to confirm the results of the present study. Finally, studies in cellular models are required to delineate the role of *RRM2B*, and other 8q-chromosome genes in cancer cell maintenance, therapeutic targeting and clinical outcomes.

## Conclusions

Overall, this study provides an in-depth analysis of *RRM2B* alterations in multiple tumor types, majorly reflected as *RRM2B* amplifications in conjunction with *MYC*, and other genes on chromosome 8. These cases exhibit a distinct 8q-amplification pattern as well as survival outcome differences and mutation signature differences, depending on cancer type and subtypes. Future genome-wide studies of other cancer datasets are warranted to confirm the results of the present study. Finally, studies in cellular models are required to delineate the role of *RRM2B*, and other 8q-chromosome genes in cancer cell maintenance, therapeutic targeting and clinical outcomes.

## Credit authorship contribution statement

**Waleed Iqbal:** Methodology, Formal Analysis, Investigation, Roles/Writing - original draft, Data Curation, Visualization, Validation. **Elena V. Demidova:** Methodology, Formal Analysis, Investigation, Roles/Writing - original draft, Writing - review & editing, Data curation, Visualization, Validation. **Samantha Serrao:** Formal Analysis, Investigation, Roles/Writing - original draft, Data curation. **Taha ValizadehAslani:** Software, Formal Analysis, Data curation, Visualization. **Gail Rosen:** Writing - review & editing, Methodology, Supervision, Funding Acquisition. **Sanjeevani Arora:** Conceptualization, Methodology, Writing - review & editing, Supervision, Project administration, Resources, Funding Acquisition.

## Acknowledgment

We are express gratitude to Dr. Erica Golemis at the Fox Chase Cancer Center for many helpful discussions and advice with this work.

## Ethics Statement

Not applicable to this study.

## Abbreviations

BLCA: Bladder urothelial carcinoma
BRCA: Breast invasive carcinoma
CESC: Cervical squamous cell carcinoma and endocervical adenocarcinoma
CNV: Copy number variations
COAD: Colon adenocarcinoma
DDR: DNA damage and response
DFS: Disease-free survival
ESCA: Esophageal carcinoma
HNSC: Head and neck squamous cell carcinoma
KIRC: Kidney renal clear cell carcinoma
LIHC: Liver hepatocellular carcinoma
LUAD: Lung adenocarcinoma
MDS: Mitochondrial depletion syndrome
MESO: Mesothelioma
NHL: Non-Hodgkin lymphoma
OV: Ovarian serous cystadenocarcinoma
OS: Overall survival
PAAD: Pancreatic adenocarcinoma
PCPG: Pheochromocytoma and paraganglioma
PRAD: Prostate adenocarcinoma
RNR: Ribonucleotide reductase
SARC: Sarcoma
SKCM: Skin cutaneous melanoma
STAD: Stomach adenocarcinoma
TMB: Tumor mutation burden
USEC: Uterine corpus endometrial carcinoma

## References

[1] Foskolou IP, Jorgensen C, Leszczynska KB, Olcina MM, Tarhonskaya H, Haisma B, D’Angiolella V, Myers WK, Domene C, Flashman E, Hammond EM. Ribonucleotide Reductase Requires Subunit Switching in Hypoxia to Maintain DNA Replication. Mol Cell 2017;66:206–20 e9.

[2] Aye Y, Li M, Long MJ, Weiss RS. Ribonucleotide reductase and cancer: biological mechanisms and targeted therapies. Oncogene 2015;34:2011–21.

[3] Okumura H, Natsugoe S, Matsumoto M, Mataki Y, Takatori H, Ishigami S, Takao S, Aikou T. The predictive value of p53, p53R2, and p21 for the effect of chemoradiation therapy on oesophageal squamous cell carcinoma. Br J Cancer 2005;92:284–9.

[4] Foskolou IP, Hammond EM. RRM2B: An oxygen-requiring protein with a role in hypoxia. Mol Cell Oncol 2017;4:e1335272.

[5] Wang X, Liu X, Xue L, Zhang K, Kuo ML, Hu S, Zhou B, Ann D, Zhang S, Yen Y. Ribonucleotide reductase subunit p53R2 regulates mitochondria homeostasis and function in KB and PC-3 cancer cells. Biochem Biophys Res Commun 2011;410:102–7.

[6] Tanaka H, Arakawa H, Yamaguchi T, Shiraishi K, Fukuda S, Matsui K, Takei Y, Nakamura Y. A ribonucleotide reductase gene involved in a p53-dependent cell-cycle checkpoint for DNA damage. Nature 2000;404:42–9.

[7] Bornstein B, Area E, Flanigan KM, Ganesh J, Jayakar P, Swoboda KJ, Coku J, Naini A, Shanske S, Tanji K, Hirano M, DiMauro S. Mitochondrial DNA depletion syndrome due to mutations in the RRM2B gene. Neuromuscul Disord 2008;18:453–9.

[8] Gorman GS, Rw T. RRM2B-Related Mitochondrial Disease. In: Adam MP, Ardinger HH, Pagon RA, Wallace SE, Bean LJH, Stephens K, Amemiya A, editors. Gene Reviews. Internet: Seattle (WA): University of Washington, Seattle, 2014.

[9] Chae YK, Anker JF, Carneiro BA, Chandra S, Kaplan J, Kalyan A, Santa-Maria CA, Platanias LC, Giles FJ. Genomic landscape of DNA repair genes in cancer. Oncotarget 2016;7:23312–21.

[10] Yanamoto S, Kawasaki G, Yoshitomi I, Mizuno A. Expression of p53R2, newly p53 target in oral normal epithelium, epithelial dysplasia and squamous cell carcinoma. Cancer Lett 2003;190:233–43.

[11] Okumura H, Natsugoe S, Yokomakura N, Kita Y, Matsumoto M, Uchikado Y, Setoyama T, Owaki T, Ishigami S, Aikou T. Expression of p53R2 is related to prognosis in patients with esophageal squamous cell carcinoma. Clin Cancer Res 2006;12:3740–5.

[12] Chen J, Xiao Y, Cai X, Liu J, Chen K, Zhang X. Overexpression of p53R2 is associated with poor prognosis in lung sarcomatoid carcinoma. BMC Cancer 2017;17:855.

[13] Liu X, Lai L, Wang X, Xue L, Leora S, Wu J, Hu S, Zhang K, Kuo ML, Zhou L, Zhang H, Wang Y, Wang Y, Zhou B, Nelson RA, Zheng S, Zhang S, Chu P, Y Y. Ribonucleotide reductase small subunit M2B prognoses better survival in colorectal cancer. Cancer Research 2011;71:3202–13.

[14] Parris TZ, Kovacs A, Hajizadeh S, Nemes S, Semaan M, Levin M, Karlsson P, Helou K. Frequent MYC coamplification and DNA hypomethylation of multiple genes on 8q in 8p11-p12-amplified breast carcinomas. Oncogenesis 2014;3:e95.

[15] Christoph F, Schmidt B, Schmitz-Drager BJ, Schulz WA. Over-expression and amplification of the c-myc gene in human urothelial carcinoma. Int J Cancer 1999;84:169–73.

[16] Kalkat M, De Melo J, Hickman KA, Lourenco C, Redel C, Resetca D, Tamachi A, Tu WB, Penn LZ. MYC Deregulation in Primary Human Cancers. Genes (Basel) 2017;8.

[17] Schleicher C, Poremba C, Wolters H, Schafer KL, Senninger N, Colombo-Benkmann M. Gain of chromosome 8q: a potential prognostic marker in resectable adenocarcinoma of the pancreas? Ann Surg Oncol 2007;14:1327–35.

[18] Kwon MJ, Kim RN, Song K, Jeon S, Jeong HM, Kim JS, Han J, Hong S, Oh E, Choi JS, An J, Pollack JR, Choi YL, Park CK, Shin YK. Genes co-amplified with ERBB2 or MET as novel potential cancer-promoting genes in gastric cancer. Oncotarget 2017;8:92209–26.

[19] Bilal E, Vassallo K, Toppmeyer D, Barnard N, Rye IH, Almendro V, Russnes H, Borresen-Dale AL, Levine AJ, Bhanot G, Ganesan S. Amplified loci on chromosomes 8 and 17 predict early relapse in ER-positive breast cancers. PLoS One 2012;7:e38575.

[20] Ehlers JP, Worley L, Onken MD, Harbour JW. DDEF1 is located in an amplified region of chromosome 8q and is overexpressed in uveal melanoma. Clin Cancer Res 2005;11:3609–13.

[21] Yong ZW, Zaini ZM, Kallarakkal TG, Karen-Ng LP, Rahman ZA, Ismail SM, Sharifah NA, Mustafa WM, Abraham MT, Tay KK, Zain RB. Genetic alterations of chromosome 8 genes in oral cancer. Sci Rep 2014;4:6073.

[22] Sato K, Qian J, Slezak JM, Lieber MM, Bostwick DG, Bergstralh EJ, Jenkins RB. Clinical significance of alterations of chromosome 8 in high-grade, advanced, nonmetastatic prostate carcinoma. J Natl Cancer Inst 1999;91:1574–80.

[23] Byrne JA, Balleine RL, Schoenberg Fejzo M, Mercieca J, Chiew YE, Livnat Y, St Heaps L, Peters GB, Byth K, Karlan BY, Slamon DJ, Harnett P, Defazio A. Tumor protein D52 (TPD52) is overexpressed and a gene amplification target in ovarian cancer. Int J Cancer 2005;117:1049–54.

[24] Ho JC, Cheung ST, Patil M, Chen X, Fan ST. Increased expression of glycosylphosphatidylinositol anchor attachment protein 1 (GPAA1) is associated with gene amplification in hepatocellular carcinoma. Int J Cancer 2006;119:1330–7.

[25] Saha S, Bardelli A, Buckhaults P, Velculescu VE, Rago C, St Croix B, Romans KE, Choti MA, Lengauer C, Kinzler KW, Vogelstein B. A phosphatase associated with metastasis of colorectal cancer. Science 2001;294:1343–6.

[26] Bruch J, Wohr G, Hautmann R, Mattfeldt T, Bruderlein S, Moller P, Sauter S, Hameister H, Vogel W, Paiss T. Chromosomal changes during progression of transitional cell carcinoma of the bladder and delineation of the amplified interval on chromosome arm 8q. Genes Chromosomes Cancer 1998;23:167–74.

[27] Smith P, Zhou B, Ho N, Yuan YC, Su L, Tsai SC, Yen Y. 2.6 A X-ray crystal structure of human p53R2, a p53-inducible ribonucleotide reductase. Biochemistry 2009;48:11134–41.

[28] Finsterer J, Zarrouk-Mahjoub S. Phenotypic and Genotypic Heterogeneity of RRM2B Variants. Neuropediatrics 2018;49:231–7.

[29] Maatta K, Rantapero T, Lindstrom A, Nykter M, Kankuri-Tammilehto M, Laasanen SL, Schleutker J. Whole-exome sequencing of Finnish hereditary breast cancer families. Eur J Hum Genet 2016;25:85–93.

[30] Kolberg M, Strand KR, Graff P, Andersson KK. Structure, function, and mechanism of ribonucleotide reductases. Biochim Biophys Acta 2004;1699:1–34.

[31] Jia Y, Chen L, Jia Q, Dou X, Xu N, Liao DJ. The well-accepted notion that gene amplification contributes to increased expression still remains, after all these years, a reasonable but unproven assumption. J Carcinog 2016;15:3.

[32] Xue L, Zhou B, Liu X, Heung Y, Chau J, Chu E, Li S, Jiang C, Un F, Yen Y. Ribonucleotide reductase small subunit p53R2 facilitates p21 induction of G1 arrest under UV irradiation. Cancer Res 2007;67:16–21.

[33] Qi JJ, Liu L, Cao JX, An GS, Li SY, Li G, Jia HT, Ni JH. E2F1 regulates p53R2 gene expression in p53-deficient cells. Mol Cell Biochem 2015;399:179–88.

[34] Cho EC, Kuo ML, Liu X, Yang L, Hsieh YC, Wang J, Cheng Y, Yen Y. Tumor suppressor FOXO3 regulates ribonucleotide reductase subunit RRM2B and impacts on survival of cancer patients. Oncotarget 2014;5:4834–44.

[35] Brunet A, Bonni A, Zigmond MJ, Lin MZ, Juo P, Hu LS, Anderson MJ, Arden KC, Blenis J, Greenberg ME. Akt promotes cell survival by phosphorylating and inhibiting a Forkhead transcription factor. Cell 1999;96:857–68.

[36] Chernov MV, Gr S. The p53 activation and apoptosis induced by DNA damage are reversibly inhibited by salicylate. Oncogene 1997;14:2503–10.

[37] Masutani M, Sonenberg N, Yokoyama S, Imataka H. Reconstitution reveals the functional core of mammalian eIF3. EMBO J 2007;26:3373–83.

[38] Morris C, Tomimatsu N, Richard DJ, Cluet D, Burma S, Khanna KK, Jalinot P. INT6/EIF3E interacts with ATM and is required for proper execution of the DNA damage response in human cells. Cancer Res 2012;72:2006–16.

[39] Koch HB, Zhang R, Verdoodt B, Bailey A, Zhang CD, Yates JR, 3rd, Menssen A, Hermeking H. Large-scale identification of c-MYC-associated proteins using a combined TAP/MudPIT approach. Cell Cycle 2007;6:205–17.

[40] Liao Y, Wang J, Jaehnig EJ, Shi Z, Zhang B. WebGestalt 2019: gene set analysis toolkit with revamped UIs and APIs. Nucleic Acids Research 2019;47:W199–W205.

[41] Campbell BB, Light N, Fabrizio D, Zatzman M, Fuligni F, de Borja R, Davidson S, Edwards M, Elvin JA, Hodel KP, Zahurancik WJ, Suo Z, Lipman T, Wimmer K, Kratz CP, Bowers DC, Laetsch TW, Dunn GP, Johanns TM, Grimmer MR, Smirnov IV, Larouche V, Samuel D, Bronsema A, Osborn M, Stearns D, Raman P, Cole KA, Storm PB, Yalon M, Opocher E, Mason G, Thomas GA, Sabel M, George B, Ziegler DS, Lindhorst S, Issai VM, Constantini S, Toledano H, Elhasid R, Farah R, Dvir R, Dirks P, Huang A, Galati MA, Chung J, Ramaswamy V, Irwin MS, Aronson M, Durno C, Taylor MD, Rechavi G, Maris JM, Bouffet E, Hawkins C, Costello JF, Meyn MS, Pursell ZF, Malkin D, Tabori U, Shlien A. Comprehensive Analysis of Hypermutation in Human Cancer. Cell 2017;171:1042–56 e10.

[42] Alexandrov LB, Kim J, Haradhvala NJ, Huang MN, Tian Ng AW, Wu Y, Boot A, Covington KR, Gordenin DA, Bergstrom EN, Islam SMA, Lopez-Bigas N, Klimczak LJ, McPherson JR, Morganella S, Sabarinathan R, Wheeler DA, Mustonen V, Group PMSW, Getz G, Rozen SG, Stratton MR, Consortium P. The repertoire of mutational signatures in human cancer. Nature 2020;578:94–101.

[43] Cadet J, Wagner JR. DNA base damage by reactive oxygen species, oxidizing agents, and UV radiation. Cold Spring Harb Perspect Biol 2013;5.

[44] El-Hattab AW, Craigen WJ, Scaglia F. Mitochondrial DNA maintenance defects. Biochim Biophys Acta Mol Basis Dis 2017;1863:1539–55.

[45] Dorman SN, Baranova K, Knoll JH, Urquhart BL, Mariani G, Carcangiu ML, Rogan PK. Genomic signatures for paclitaxel and gemcitabine resistance in breast cancer derived by machine learning. Mol Oncol 2016;10:85–100.

[46] Gatto F, Miess H, Schulze A, Nielsen J. Flux balance analysis predicts essential genes in clear cell renal cell carcinoma metabolism. Sci Rep 2015;5:10738.

[47] Macintyre G, Goranova TE, De Silva D, Ennis D, Piskorz AM, Eldridge M, Sie D, Lewsley LA, Hanif A, Wilson C, Dowson S, Glasspool RM, Lockley M, Brockbank E, Montes A, Walther A, Sundar S, Edmondson R, Hall GD, Clamp A, Gourley C, Hall M, Fotopoulou C, Gabra H, Paul J, Supernat A, Millan D, Hoyle A, Bryson G, Nourse C, Mincarelli L, Sanchez LN, Ylstra B, Jimenez-Linan M, Moore L, Hofmann O, Markowetz F, McNeish IA, Brenton JD. Copy number signatures and mutational processes in ovarian carcinoma. Nat Genet 2018;50:1262–70.

[48] Ramón YCS, Sesé M, Capdevila C, Aasen T, De Mattos-Arruda L, Diaz-Cano SJ, Hernández-Losa J, Castellví J. Clinical implications of intratumor heterogeneity: challenges and opportunities. Journal of molecular medicine (Berlin, Germany) 2020;98:161–77.

[49] Mirzayans R, Murray D. Intratumor Heterogeneity and Therapy Resistance: Contributions of Dormancy, Apoptosis Reversal (Anastasis) and Cell Fusion to Disease Recurrence. International journal of molecular sciences 2020;21.

[50] Ballabio S, Craparotta I, Paracchini L, Mannarino L, Corso S, Pezzotta MG, Vescio M, Fruscio R, Romualdi C, Dainese E, Ceppi L, Calura E, Pileggi S, Siravegna G, Pattini L, Martini P, Delle Marchette M, Mangioni C, Ardizzoia A, Pellegrino A, Landoni F, D’Incalci M, Beltrame L, Marchini S. Multisite analysis of high-grade serous epithelial ovarian cancers identifies genomic regions of focal and recurrent copy number alteration in 3q26.2 and 8q24.3. Int J Cancer 2019;145:2670–81.

[51] Luo J, Solimini NL, Elledge SJ. Principles of cancer therapy: oncogene and non-oncogene addiction. Cell 2009;136:823–37.

[52] Solimini NL, Luo J, Elledge SJ. Non-oncogene addiction and the stress phenotype of cancer cells. Cell 2007;130:986–8.

[53] Steinberga I, Jansson K, Sorbe B. Quality Indicators and Survival Outcome in Stage IIIB-IVB Epithelial Ovarian Cancer Treated at a Single Institution. In vivo (Athens, Greece) 2019;33:1521–30.

[54] Xu X, Page JL, Surtees JA, Liu H, Lagedrost S, Lu Y, Bronson R, Alani E, Nikitin AY, Weiss RS. Broad overexpression of ribonucleotide reductase genes in mice specifically induces lung neoplasms. Cancer Res 2008;68:2652–60.

[55] Kuo ML, Sy AJ, Xue L, Chi M, Lee MT, Yen T, Chiang MI, Chang L, Chu P, Yen Y. RRM2B suppresses activation of the oxidative stress pathway and is up-regulated by p53 during senescence. Sci Rep 2012;2:822.

[56] Desai A, Yan Y, Gerson SL. Advances in therapeutic targeting of the DNA damage response in cancer. DNA Repair (Amst) 2018;66-67:24–9.

[57] Nickoloff JA, Jones D, Lee SH, Williamson EA, Hromas R. Drugging the Cancers Addicted to DNA Repair. J Natl Cancer Inst 2017;109.

[58] Ma J, Setton J, Lee NY, Riaz N, Powell SN. The therapeutic significance of mutational signatures from DNA repair deficiency in cancer. Nature Communications 2018;9:3292.

[59] O’Connor MJ. Targeting the DNA Damage Response in Cancer. Mol Cell 2015;60:547–60.

[60] Cerami E, Gao J, Dogrusoz U, Gross BE, Sumer SO, Aksoy BA, Jacobsen A, Byrne CJ, Heuer ML, Larsson E, Antipin Y, Reva B, Goldberg AP, Sander C, Schultz N. The cBio cancer genomics portal: an open platform for exploring multidimensional cancer genomics data. Cancer Discov 2012;2:401–4.

[61] Fisher RA. On the Interpretation of χ^2^ from Contingency Tables, and the Calculation of P. Journal of the Royal Statistical Society 1922;85:87–94.

[62] Li J, Tibshirani R. Finding consistent patterns: a nonparametric approach for identifying differential expression in RNA-Seq data. Stat Methods Med Res 2013;22:519–36.

[63] Gao X, Song PX. Nonparametric tests for differential gene expression and interaction effects in multi-factorial microarray experiments. BMC Bioinformatics 2005;6:186.

[64] Karl P, Francis G. Note on regression and inheritance in the case of two parents. Proc R Soc Lond 1997;58:240–2.

[65] Mantel N. Evaluation of survival data and two new rank order statistics arising in its consideration. Cancer chemotherapy reports 1966;50:163–70.

[66] Experimental Design: Procedures for the Behavioral Sciences. Thousand Oaks, California, 2013.

[67] Shuster JJ, Theriaque DW, Ilfeld BM. Applying Hodges-Lehmann scale parameter estimates to hospital discharge times. Clin Trials 2008;5:631–4.

[68] Fujikoshi Y. Two-way ANOVA models with unbalanced data. Discrete Mathematics 1993;116:315–34.

[69] Benjamini Y, Hochberg Y. Controlling the False Discovery Rate: A Practical and Powerful Approach to Multiple Testing. 1995;57:289–300.

[70] Kaplan EL, Meier P. Nonparametric Estimation from Incomplete Observations. Journal of the American Statistical Association 1958;53:457–81.

[71] Hazra A, Gogtay N. Biostatistics Series Module 9: Survival Analysis. Indian journal of dermatology 2017;62:251–7.

[72] Oughtred R, Stark C, Breitkreutz BJ, Rust J, Boucher L, Chang C, Kolas N, O’Donnell L, Leung G, McAdam R, Zhang F, Dolma S, Willems A, Coulombe-Huntington J, Chatr-Aryamontri A, Dolinski K, Tyers M. The BioGRID interaction database: 2019 update. Nucleic Acids Res 2019;47:D529–D41.

[73] Shannon P, Markiel A, Ozier O, Baliga NS, Wang JT, Ramage D, Amin N, Schwikowski B, Ideker T. Cytoscape: a software environment for integrated models of biomolecular interaction networks. Genome Res 2003;13:2498–504.

[74] Tebbi A, Guittet O, Tuphile K, Cabrie A, Lepoivre M. Caspase-dependent Proteolysis of Human Ribonucleotide Reductase Small Subunits R2 and p53R2 during Apoptosis. J Biol Chem 2015;290:14077–90.

[75] Piao C, Youn CK, Jin M, Yoon SP, Chang IY, Lee JH, You HJ. MEK2 regulates ribonucleotide reductase activity through functional interaction with ribonucleotide reductase small subunit p53R2. Cell Cycle 2012;11:3237–49.

