## Supplementary figures for "*RRM2B* is frequently amplified across multiple tumor types: non-oncogenic addiction and therapeutic opportunities"

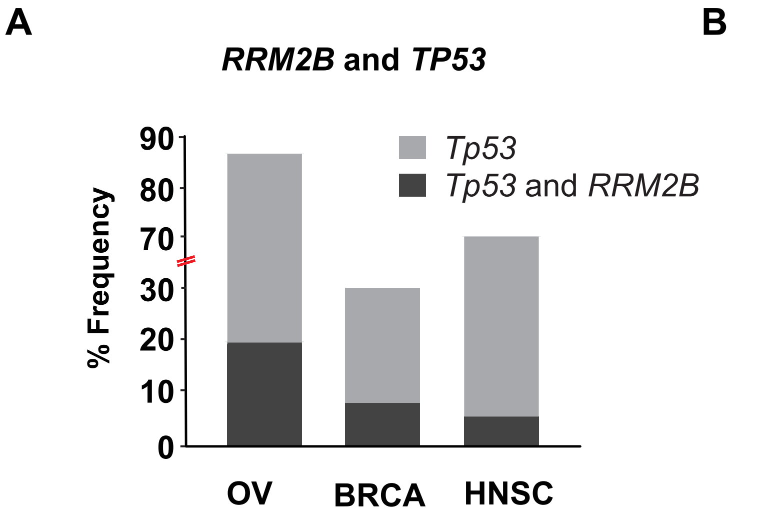


Iqbal, Demidova et al., Supplementary Figure 1

#### Ovarian Cancer (OV)

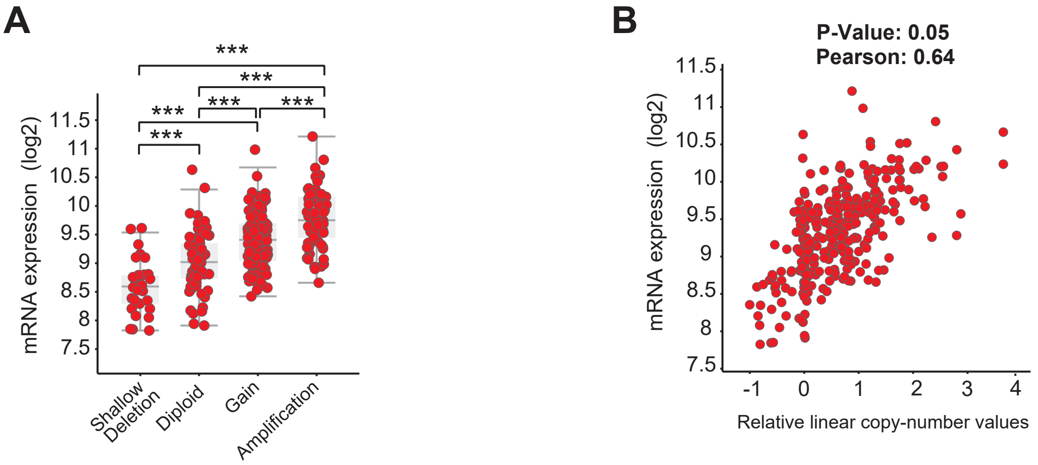


#### Breast Cancer (BRCA)

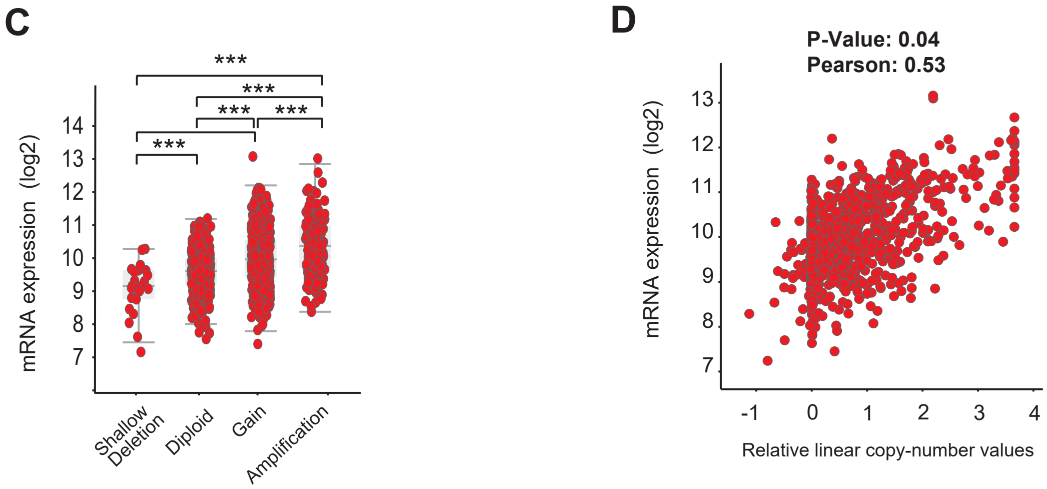


#### Head and Neck Cancer (HNSC)

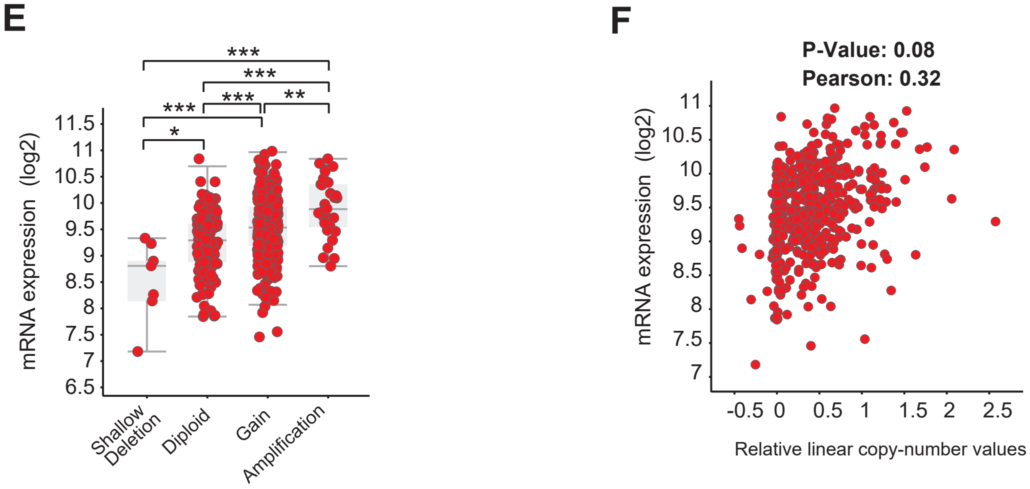


**Iqbal, Demidova et al., Supplementary Figure 2**

### Breast Cancer (BRCA)

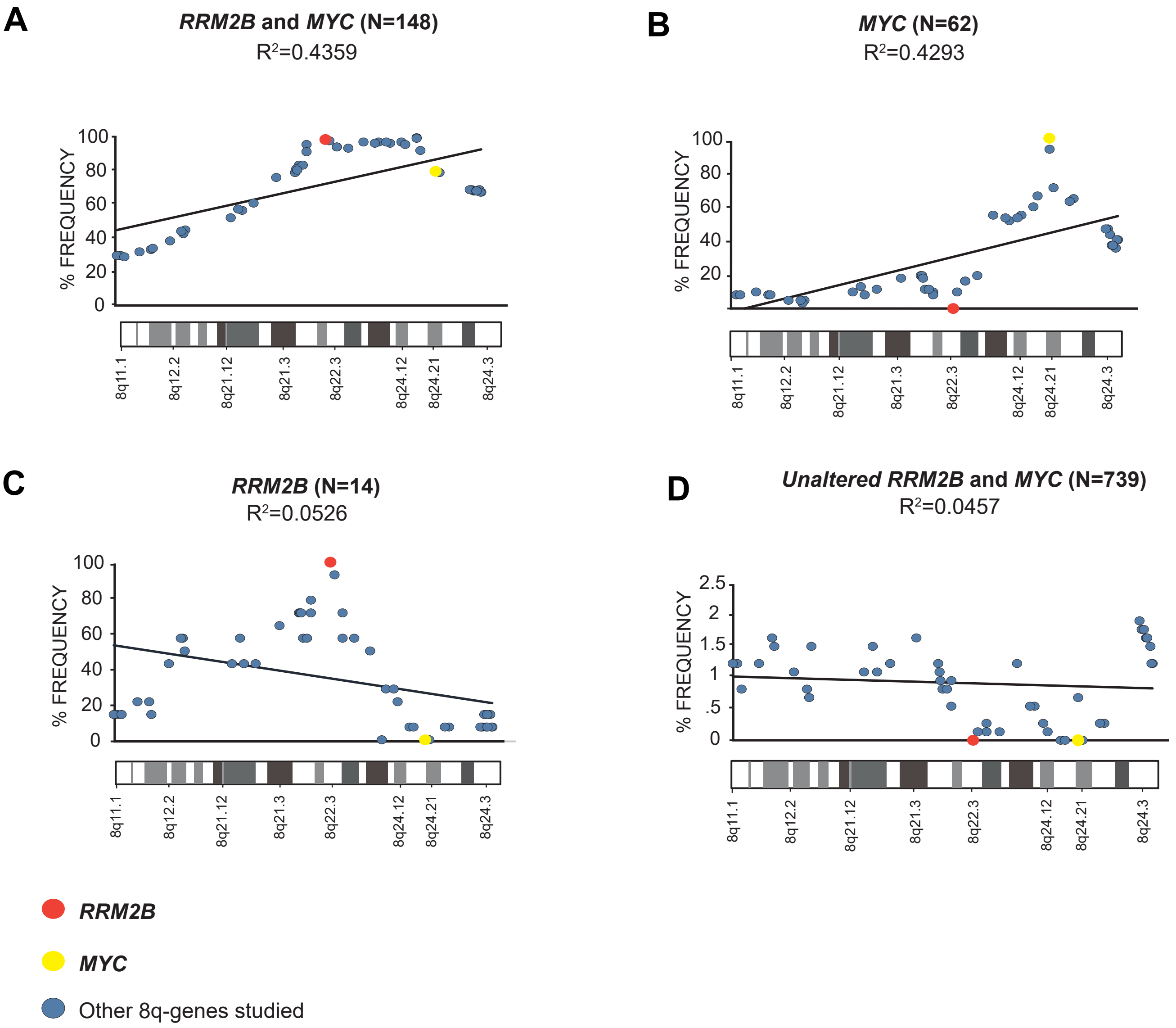


Iqbal, Demidova et al., Supplementary Figure 3

#### Head and Neck Cancer (HNSC)

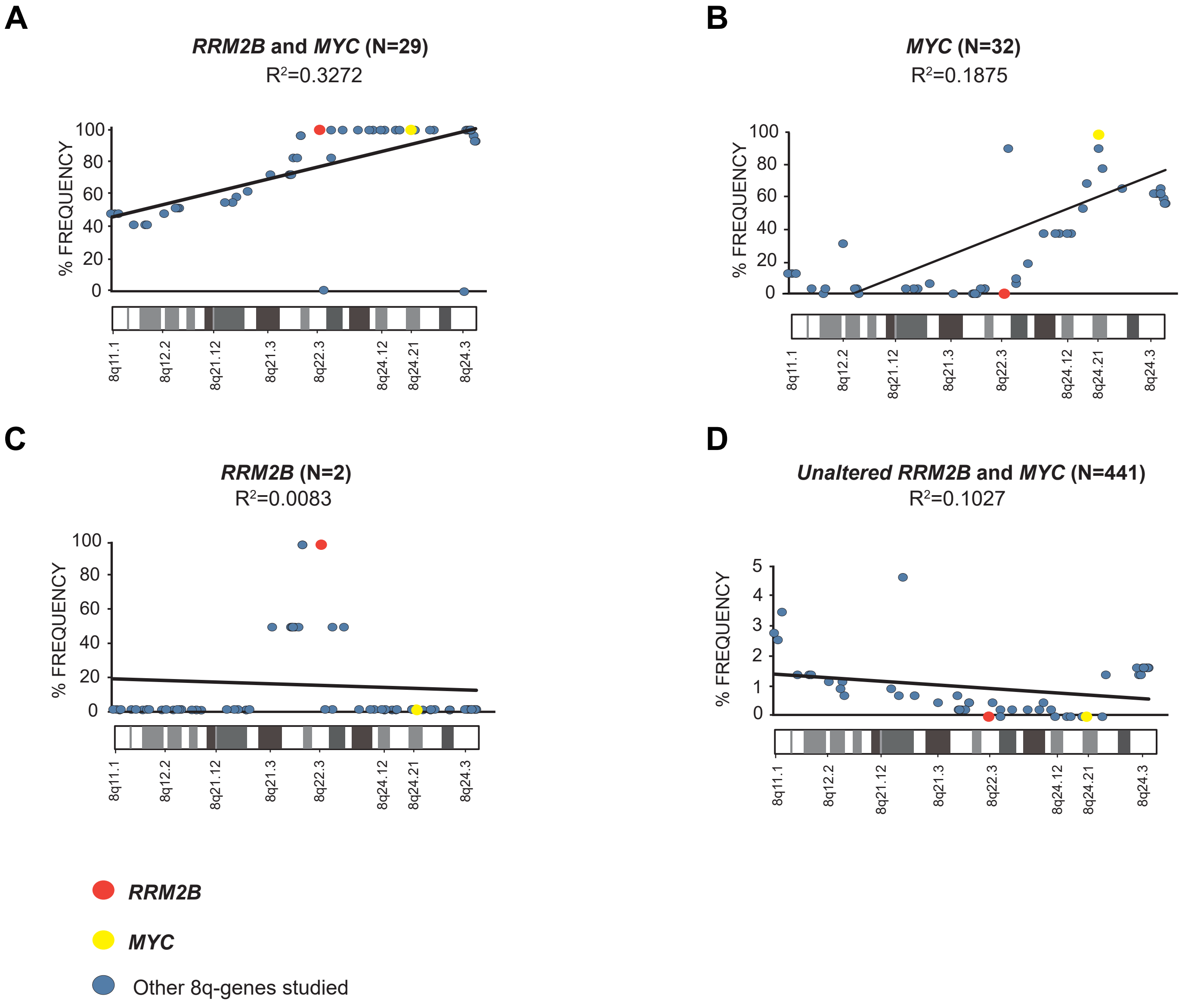


Iqbal, Demidova et al., Supplementary Figure 4

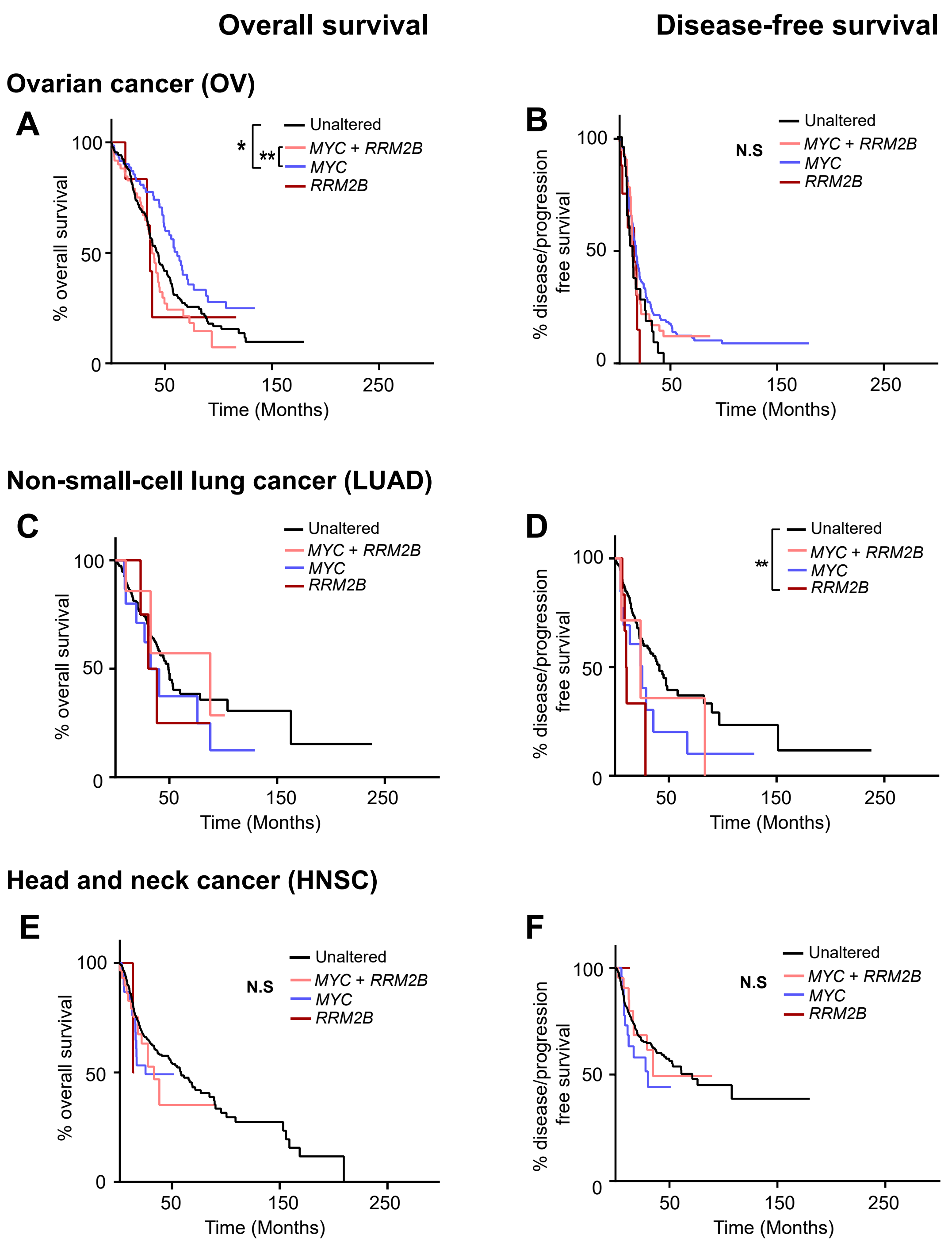


**Iqbal, Demidova et al., Supplementary Figure 5**

#### Ovarian cancer (OV)

**A**

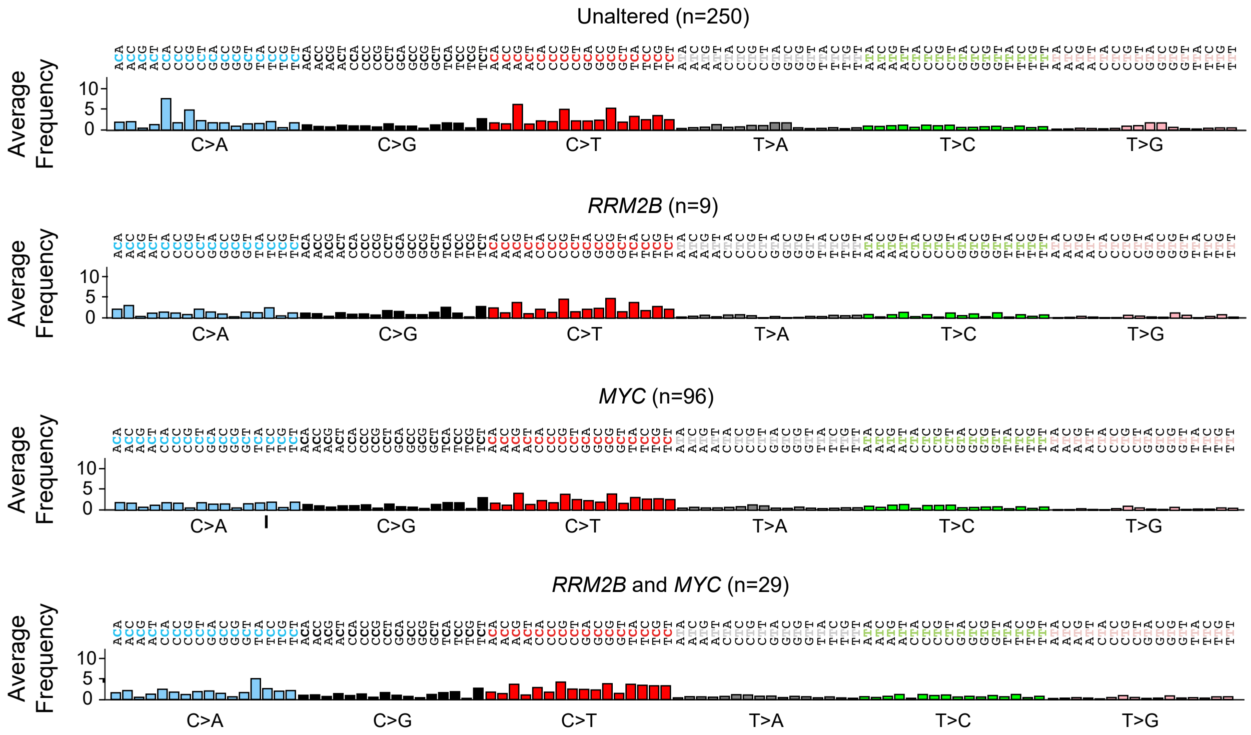


#### Head and Neck cancer (HNSC)

**B**

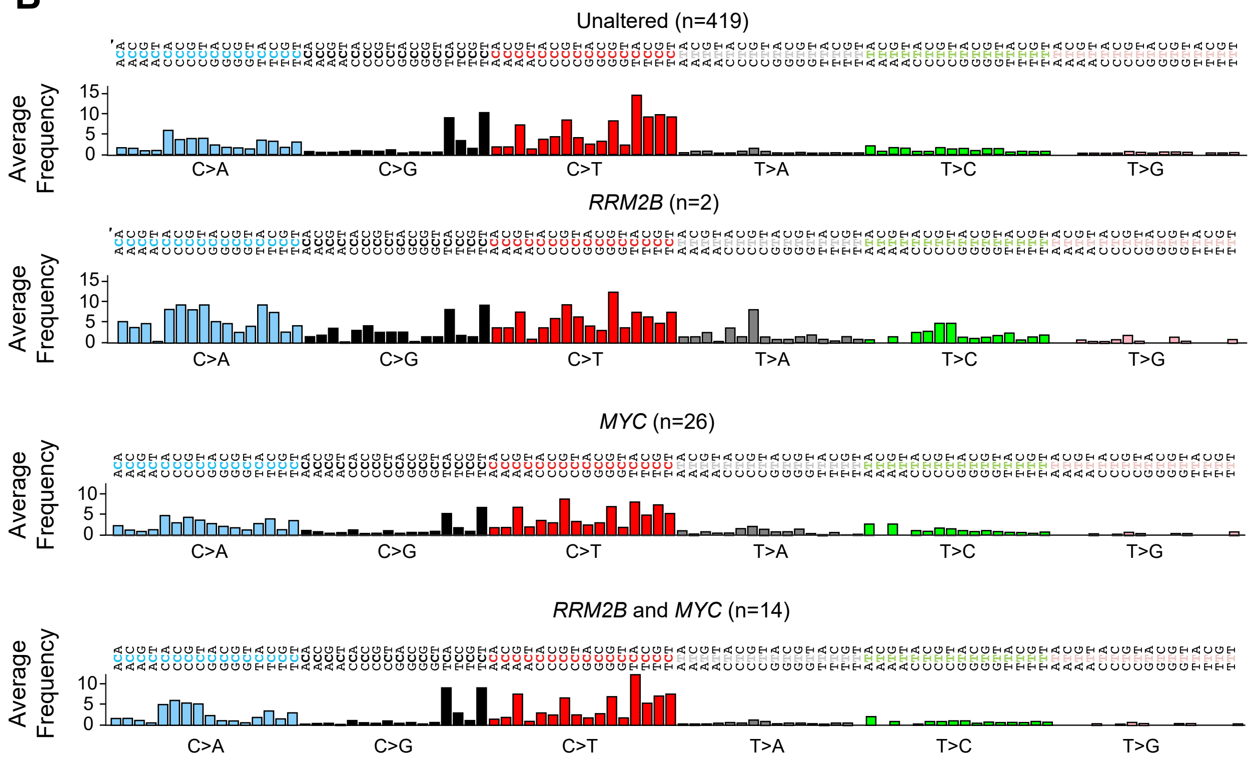


**Iqbal, Demidova et al, Supplementary Figure 6**

C>A C>G C>T T>A T>C T>G

#### Non-small cell lung cancer (LUAD)

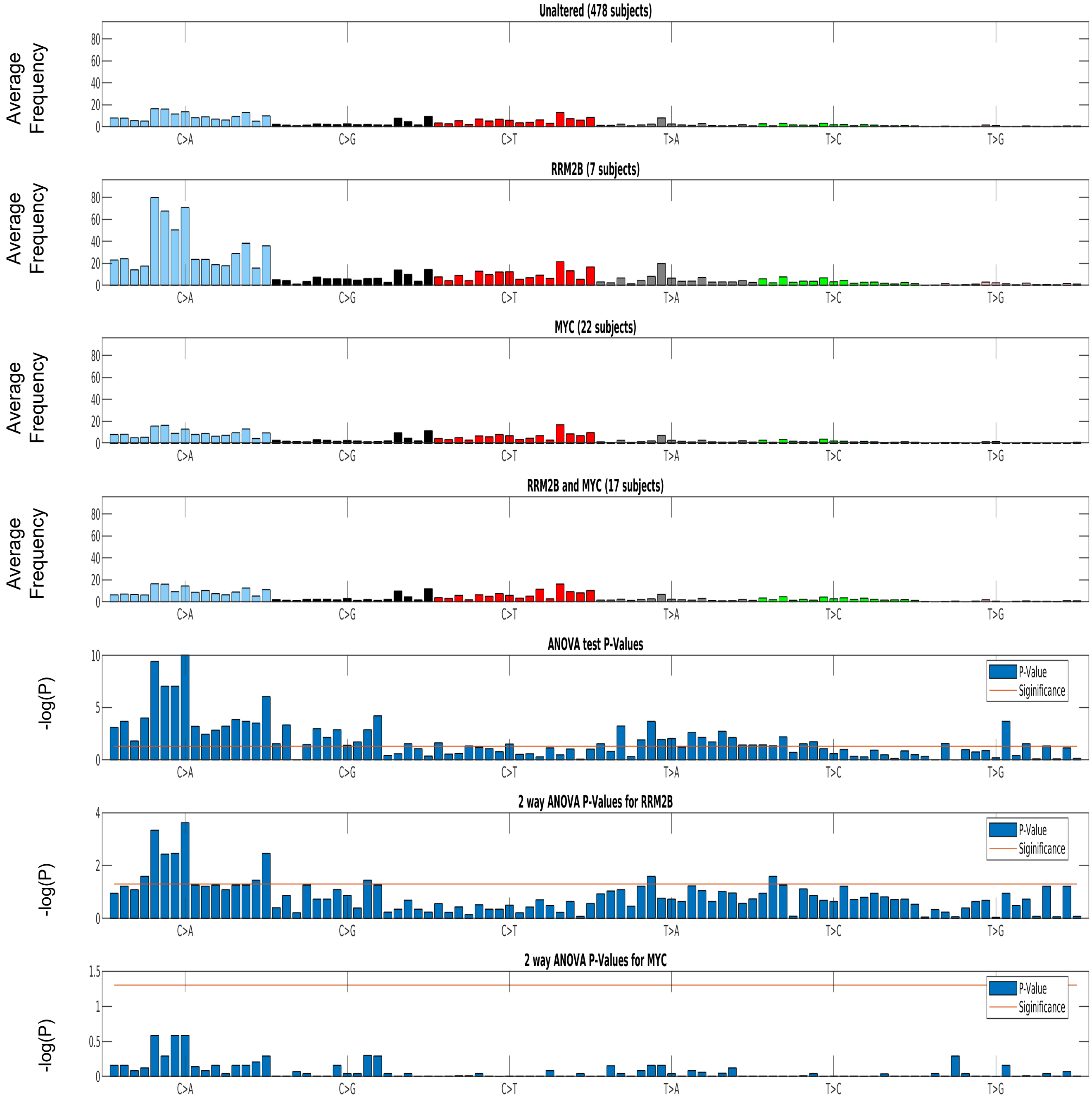


#### Iqbal, Demidova et al, Supplementary Figure 7
