## Supplementary legends for "*RRM2B* is frequently amplified across multiple tumor types: non-oncogenic addiction and therapeutic opportunities"

**Supplementary figures legends.**

**Supplementary Figure 1.** Somatic alteration frequencies in RRM2B along with either MYC or Tp53 in OV, BRCA and HNSC. A: Comparison of frequency between cases with both RRM2B and Tp53 alterations (dark gray) and cases with Tp53 alterations alone (light gray). B: Comparison of frequency between cases with both RRM2B and MYC alterations (dark gray) and cases with MYC alterations alone (light gray).

**Supplementary figure 2.** *RRM2B* mRNA expression based on copy number alterations in ovarian (OV), breast (BRCA) and head and neck (HNSC) cancers. A, C, E: log2 mRNA expression values from cases with different copy number alterations (shallow deletion to amplifications). Mann-Whitney Non-Parametric T-tests were performed on the data and differences between the alteration types is represented by P-value *<0.05, ** for P-value <0.01 and *** for P-value <0.001. B, D, F: log2 mRNA expression values based on relative copy number alterations in *RRM2B*. The Spearman correlation values (for correlation between increase in mRNA expression and increase in relative copy number values) are listed for each comparison.

**Supplementary Figure 3.** Amplification frequencies of 8q-genes in breast cancer (BRCA) study. A: cases with co-amplification of RRM2B and MYC. B: cases MYC only amplification. C: cases RRM2B only amplifications. D: cases with neither (unaltered) were plotted as percent of frequency for amplifications in various 8q-region genes relevant for cancer (see Supplementary Table 1). RRM2B (red circle), MYC (yellow circle), and other genes (blue circle). The Pearson correlation (R^2^ value) for the data points is represented by a black trend line.

**Supplementary Figure 4.** Amplification frequencies of 8q-genes in head and neck cancer (HNSC) study. A: cases with co-amplification of RRM2B and MYC. B: cases MYC only amplification. C: cases RRM2B only amplifications. D: cases with neither (unaltered) were plotted as percent of frequency for amplifications in various 8q-region genes relevant for cancer (see Supplementary Table 1). RRM2B (red circle), MYC (yellow circle), and other genes (blue circle). The Pearson correlation (R^2^ value) for the data points is represented by a black trend line.

**Supplementary figure 5.** Kaplan-Meier curves for overall survival (OS) and disease/progression-free survival (DFS) in ovarian cancer (A, B), non-small cell lung cancer (C, D) and head and neck cancer (E, F) studies with *RRM2B* amplifications and/or *MYC* amplifications, and those that are unaltered for *RRM2B* and *MYC*. A: OS in OV cases. The cases that were unaltered for both genes (black, n=174), cases with *RRM2B* amplifications (red, n=8), cases with *MYC* amplifications (blue, n=72) and co-amplifications (pink, n=60) were plotted. B: DFS in OV cases. The cases that were unaltered for both genes (black, n=144), cases with *RRM2B* amplifications (red, n=7), cases with *MYC* amplifications (blue, n=65) and co-amplifications (pink, n= 52) were plotted. C: OS in LUAD cases. The cases that were unaltered for both genes (black, n=195), cases with *RRM2B* amplifications (red, n=6), cases with *MYC* amplifications (blue, n=15) and co-amplifications (pink, n=7) were plotted. D: DFS in LUAD cases. The cases that were unaltered for both genes (black, n=162), cases with *RRM2B* amplifications (red, n=6), cases with *MYC* amplifications (blue, n=13) and cases with amplifications in both (pink, n=7) were plotted. E: OS in HNSC cases. The cases that were unaltered for both genes (black, n=431), cases with *RRM2B* amplifications (red, n=2), cases with *MYC* amplifications (blue, n=32) and co-amplifications (pink, n=30) were plotted. F: DFS in HNSC cases. The cases that were unaltered for both genes (black, n=327), cases with *RRM2B* amplifications (red, n=1), cases with *MYC* amplifications (blue, n=25) and co-amplifications (pink, n=327) were plotted. The plots were compared using Log-rank test and significance is shown as follows: P-value *<0.05, ** P-value <0.01 and ***P-value <0.001.

**Supplementary Figure 6.** Mutation signatures of OV and HNSC cancers based on RRM2B or MYC amplifications. Tumor whole-exome sequence data from the PanCancer Atlas studies was used to calculate the average frequency of the 96 trinucleotide context mutations in each group: unaltered cases, cases with RRM2B or MYC amplifications only, and cases with both. A: Mutation Signatures in OV cancer. B: Mutation Signatures in HNSC.

**Supplementary Figure *7*.** Mutation signature of lung cancer (LUAD) patients based on *RRM2B* or *MYC* amplifications. A one-way ANOVA (*RRM2B* amplifications only versus other groups) and two-way ANOVA (included group with *RRM2B* and *MYC* co-amplifications) analysis showed significant C>A mutations. Top: Tumor whole-exome sequence data from the PanCancer Atlas studies was used to calculate the average frequency of the 96 trinucleotide context mutations in each group: unaltered cases, cases with *RRM2B* or *MYC* amplifications only, and cases with both. Bottom: The statistical significance of each comparison is represented by an inverse transformed p-value, -log_10_ (P-value), calculated by an ANOVA test on each group of signatures (96 trinucleotide context mutations) compared. The -log_10_ (P) results are provided for: one-way ANOVA comparing the *RRM2B*-only group to the other groups (panel 1) and a two-way ANOVA comparing all groups with *RRM2B* (panel 2) or *MYC* amplifications (panel 3). Since -log10 (P) is employed here, longer bars correspond to smaller P-Values, with bars above the red line being P-values less than 0.05. P-values of each subfigure have been corrected using Benjamini–Hochberg procedure.

**Supplementary tables legends.**

**Supplementary Table 1.** 8q gene list and associated Gene Ontology terms. The table on the left represents a list of the 8q-located genes (8q11-8q24) with their positions. On the right is Gene Ontology terms for the 8q-genes of interest.

**Supplementary table 2.** mRNA expression based on copy number alterations in ovarian (OV), head and neck (HNSC), and breast (BRCA) cancers of 8q amplicon genes (*CCNE2*, *EI3FE, MTDH, MYC, RAD21, TP53INP1, YWHAZ*). Mean and median mRNA expression values from cases with different copy number alterations (shallow deletion to amplifications) are shown. Mann-Whitney Non-Parametric T-tests were performed on the data and differences between the alteration types is represented by P-values.

**Supplementary Table 3.** Enrichment of specific pathways based on genes of interest. WebGestalt tool was used to perform over representation analysis, and the results were prioritized based on p-values and FDR thresholds at 0.01.

**Supplementary table 4.** Kaplan-Meier curves statistics. P-values for Kaplan-Meier curves, presented in Figure 5 and Supplementary Figure 5 for the cases with *RRM2B* amplifications and/or *MYC* amplifications, and those that are unaltered for *RRM2B* and *MYC*. The data was obtained using Log-rank test. OS, Overall Survival, DFS, Disease-Free Progression.

**Supplementary Table 5.** Tumor Mutation Burden (TMB) profile in OV, BRCA and HNSC. The data is presented for TCGA studies cohorts on ovarian, breast and head and neck cases (OV, BRCA, and HNSC correspondingly). Average TMB data shown for patients with or without *RRM2B* amplifications separately.

**Supplementary Table 6.** Mutation Signature statistics for BRCA and LUAD. The 96-trinucleotide mutation signature context statistics are represented for breast (BRCA) and lung (LUAD) TCGA cancer cohorts. One-way (*RRM2B* amplifications only versus other groups), two-way ANOVA (included group with *RRM2B* and *MYC* co-amplifications), and Wilcoxon rank-sum tests have been performed. Benjamini–Hochberg procedure has been applied to results of each column to avoid false discovery of multiple comparison. Significant p-values are highlighted in bold (p<0.05).
