## Supplementary tables for "*RRM2B* is frequently amplified across multiple tumor types: non-oncogenic addiction and therapeutic opportunities"

**Supplementary Table 1.** 8q gene list and associated Gene Ontology terms.

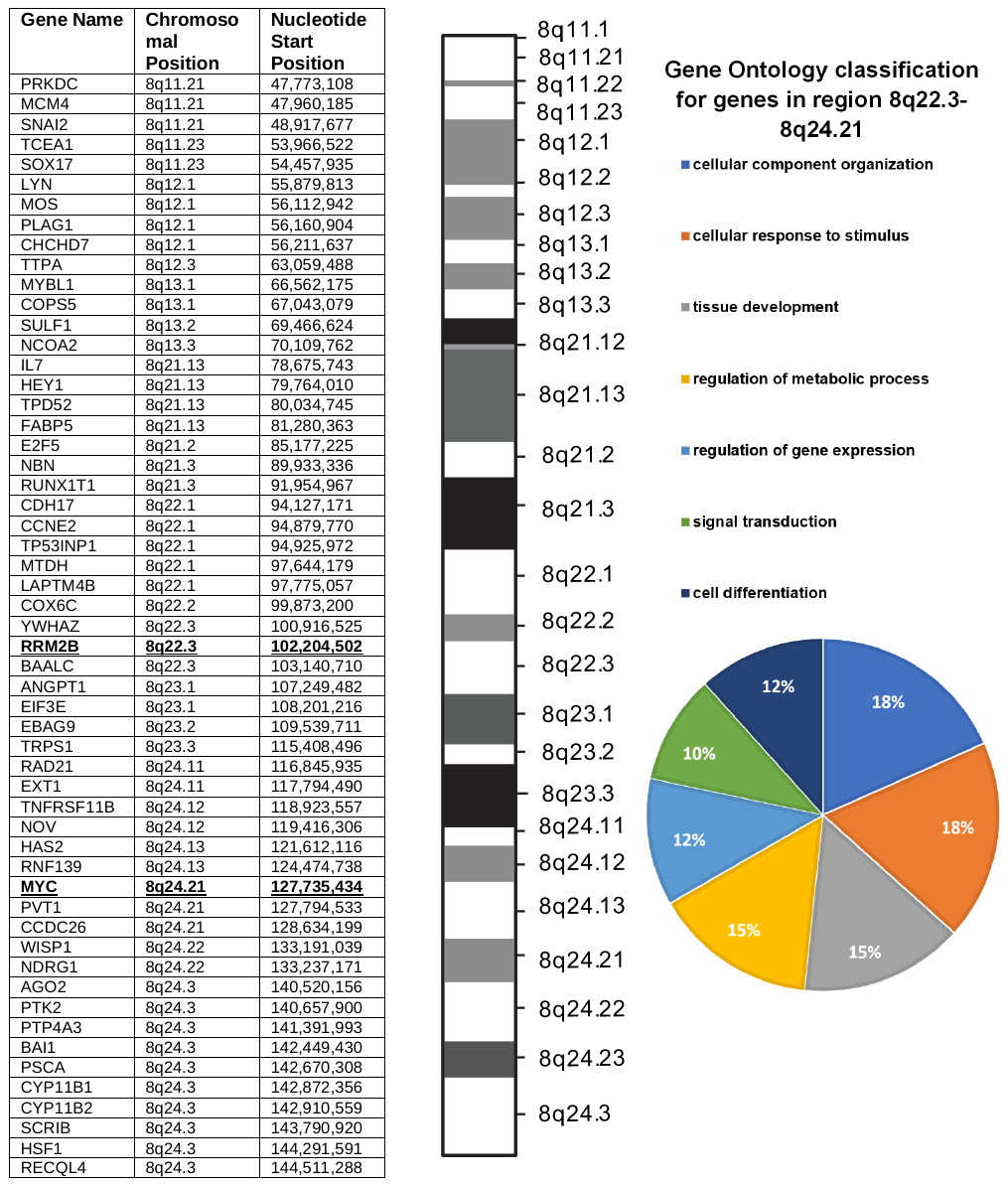

**Supplementary table 2.** mRNA expression based on copy number alterations in ovarian (OV), head and neck (HNSC), and breast (BRCA) cancers of 8q amplicon genes (*CCNE2*, *EI3FE, MTDH, MYC, RAD21, TP53INP1, YWHAZ*).

| **Ovarian cancer** | | | | | |
| --- | --- | --- | --- | --- | --- |
| **Gene Name** | **CNA Type** | **Number of Cases** | **Mean RNA expression** | **Median RNA expression** | **Standard Deviation** |
| *CCNE2* | Shallow Deletion | 26 | 123.2440324 | 87.1696835 | 116.7281103 |
|  | Diploid | 45 | 209.9187637 | 113.312714 | 256.0795475 |
|  | Gain | 120 | 192.5782797 | 143.381862 | 145.4034984 |
|  | Amplification | 10 | 188.1584144 | 156.716141 | 109.8653981 |
| *EI3FE* | Shallow Deletion | 16 | 4322.626424 | 4202.170021 | 1080.777821 |
|  | Diploid | 41 | 7860.928264 | 6130.690738 | 3786.006383 |
|  | Gain | 107 | 10293.85155 | 8552.470596 | 5056.509654 |
|  | Amplification | 36 | 13724.90475 | 11822.69819 | 6108.487215 |
| *MTDH* | Shallow Deletion | 23 | 1985.483477 | 1839.538801 | 505.3383572 |
|  | Diploid | 42 | 3043.223448 | 3119.32204 | 520.9083051 |
|  | Gain | 123 | 4283.153188 | 4046.343072 | 1242.220732 |
|  | Amplification | 12 | 6065.734514 | 5729.204805 | 1285.296511 |
| *MYC* | Shallow Deletion | 6 | 1613.205991 | 1467.957074 | 1055.766322 |
|  | Diploid | 25 | 2523.209107 | 2199.243357 | 1722.204053 |
|  | Gain | 99 | 3265.72128 | 2576.718463 | 2862.849079 |
|  | Amplification | 71 | 4270.833232 | 3526.916276 | 3014.227893 |
| *RAD21* | Shallow Deletion | 12 | 4101.070243 | 3977.527492 | 1191.775697 |
|  | Diploid | 37 | 5575.767866 | 5472.398702 | 1398.28967 |
|  | Gain | 103 | 8108.534708 | 7603.620902 | 2763.755896 |
|  | Amplification | 47 | 10526.77434 | 10016.29332 | 3384.338825 |
| *TP53INP1* | Shallow Deletion | 25 | 678.2348089 | 649.521302 | 328.1994729 |
|  | Diploid | 45 | 947.5185295 | 849.699119 | 565.0942353 |
|  | Gain | 120 | 1148.187625 | 1060.517722 | 533.9033314 |
|  | Amplification | 11 | 1223.366429 | 1261.829313 | 381.2496199 |
| *YWHAZ* | Shallow Deletion | 19 | 12878.69721 | 12057.38649 | 4534.646689 |
|  | Diploid | 42 | 18152.99435 | 17426.7053 | 4367.063183 |
|  | Gain | 117 | 26987.68132 | 25889.44422 | 8930.674452 |
|  | Amplification | 23 | 40773.0947 | 34910.24061 | 19211.74257 |
| **Gene Name** | **CNA TYPE** | **CNA TYPE COMPARED (P-Values)** | | | |
| *CCNE2* |  |  |  |  |  |
|  |  | Diploid | Gain | Amplification |  |
|  | Shallow Deletion | **0.0315** | **0.0004** | **0.0078** |  |
|  | Diploid |  | 0.1701 | 0.2563 |  |
|  | Gain |  |  | 0.6825 |  |
| *EI3FE* |  |  |  |  |  |
|  |  | Diploid | Gain | Amplification |  |
|  | Shallow Deletion | **<0.0001** | **<0.0001** | **<0.0001** |  |
|  | Diploid |  | **0.003** | **<0.0001** |  |
|  | Gain |  |  | **0.0002** |  |
| *MTDH* |  |  |  |  |  |
|  |  | Diploid | Gain | Amplification |  |
|  | Shallow Deletion | **<0.0001** | **<0.0001** | **<0.0001** |  |
|  | Diploid |  | **<0.0001** | **<0.0001** |  |
|  | Gain |  |  | **<0.0001** |  |
| *MYC* |  |  |  |  |  |
|  |  | Diploid | Gain | Amplification |  |
|  | Shallow Deletion | 0.728 | 0.1866 | 0.0762 |  |
|  | Diploid |  | **<0.0001** | **<0.0001** |  |
|  | Gain |  |  | **0.0157** |  |
| *RAD21* |  |  |  |  |  |
|  |  | Diploid | Gain | Amplification |  |
|  | Shallow Deletion | **0.02** | **0.0024** | **0.0031** |  |
|  | Diploid |  | **<0.0001** | **<0.0001** |  |
|  | Gain |  |  | **<0.0001** |  |
| *TP53INP1* |  |  |  |  |  |
|  |  | Diploid | Gain | Amplification |  |
|  | Shallow Deletion | 0.4539 | 0.6819 | 0.7675 |  |
|  | Diploid |  | 0.2034 | 0.6725 |  |
|  | Gain |  |  | 0.9392 |  |
| *YWHAZ* |  |  |  |  |  |
|  |  | Diploid | Gain | Amplification |  |
|  | Shallow Deletion | **0.0021** | **<0.0001** | **<0.0001** |  |
|  | Diploid |  | **<0.0001** | **<0.0001** |  |

| **Head and Neck Cancer** | | | | | |
| --- | --- | --- | --- | --- | --- |
| **Gene Name** | **CNA Type** | **Number of Cases** | **Mean RNA expression** | **Median RNA expression** | **Standard Deviation** |
| *CCNE2* | Shallow Deletion | 12 | 223.2757667 | 124.1725 | 256.9787484 |
|  | Diploid | 151 | 263.9824318 | 184.859 | 193.7587002 |
|  | Gain | 311 | 274.6949794 | 205.767 | 392.7472116 |
|  | Amplification | 14 | 340.2525 | 324.4195 | 134.33188 |
| *EI3FE* | Shallow Deletion | 5 | 2819.746 | 2581.26 | 659.5720811 |
|  | Diploid | 138 | 4471.594783 | 4157.4 | 1409.801988 |
|  | Gain | 324 | 6237.617191 | 5715.34 | 2428.541576 |
|  | Amplification | 21 | 8041.957619 | 7555 | 2368.236253 |
| *MTDH* | Shallow Deletion | 11 | 2568.683636 | 2542.79 | 453.2158519 |
|  | Diploid | 149 | 3395.265973 | 3397.28 | 932.9099973 |
|  | Gain | 313 | 4735.933099 | 4613.79 | 1265.39875 |
|  | Amplification | 15 | 5544.302 | 5843.11 | 884.9874173 |
| *MYC* | Shallow Deletion | 4 | 3514.535 | 2838.17 | 1991.987904 |
|  | Diploid | 120 | 4002.405333 | 3435.2 | 2489.635662 |
|  | Gain | 318 | 5078.380189 | 4621.595 | 2591.221982 |
|  | Amplification | 46 | 6354.020652 | 5129.045 | 3529.150119 |
| *RAD21* | Shallow Deletion | 3 | 3111.743333 | 3279.88 | 533.6869657 |
|  | Diploid | 127 | 4695.996693 | 4524.83 | 1344.311437 |
|  | Gain | 330 | 5954.440455 | 5717.125 | 2051.913162 |
|  | Amplification | 28 | 8196.904643 | 8299.095 | 2079.876466 |
| *TP53INP1* | Shallow Deletion | 11 | 698.7549091 | 642.27 | 503.3153836 |
|  | Diploid | 152 | 729.7924895 | 664.0755 | 367.4829155 |
|  | Gain | 311 | 694.4287678 | 629.521 | 372.3898752 |
|  | Amplification | 14 | 691.8885714 | 611.233 | 355.0717748 |
| *YWHAZ* | Shallow Deletion | 8 | 17769.5625 | 16549.9 | 4705.175738 |
|  | Diploid | 144 | 27722.85903 | 24917.55 | 10101.68784 |
|  | Gain | 315 | 41253.54073 | 39459 | 15222.38768 |
|  | Amplification | 21 | 60129.28571 | 52264.4 | 28023.13264 |
| **Gene Name** | **CNA TYPE** | **CNA TYPE COMPARED (P-Values)** | | | |
| *CCNE2* |  |  |  |  |  |
|  |  | Diploid | Gain | Amplification |  |
|  | Shallow Deletion | 0.0893 | **0.0472** | **0.0077** |  |
|  | Diploid |  | 0.7225 | **0.0147** |  |
|  | Gain |  |  | **0.0046** |  |
| *EI3FE* |  |  |  |  |  |
|  |  | Diploid | Gain | Amplification |  |
|  | Shallow Deletion | **0.0022** | **<0.0001** | **<0.0001** |  |
|  | Diploid |  | **<0.0001** | **<0.0001** |  |
|  | Gain |  |  | **<0.0001** |  |
| *MTDH* |  |  |  |  |  |
|  |  | Diploid | Gain | Amplification |  |
|  | Shallow Deletion | **0.0018** | **<0.0001** | **<0.0001** |  |
|  | Diploid |  | **<0.0001** | **<0.0001** |  |
|  | Gain |  |  | **0.0029** |  |
| *MYC* |  |  |  |  |  |
|  |  | Diploid | Gain | Amplification |  |
|  | Shallow Deletion | 0.728 | 0.1866 | 0.0762 |  |
|  | Diploid |  | **<0.0001** | **<0.0001** |  |
|  | Gain |  |  | **0.0157** |  |
| *RAD21* |  |  |  |  |  |
|  |  | Diploid | Gain | Amplification |  |
|  | Shallow Deletion | **0.02** | **0.0024** | **0.0031** |  |
|  | Diploid |  | **<0.0001** | **<0.0001** |  |
|  | Gain |  |  | **<0.0001** |  |
| *TP53INP1* |  |  |  |  |  |
|  |  | Diploid | Gain | Amplification |  |
|  | Shallow Deletion | 0.4539 | 0.6819 | 0.7675 |  |
|  | Diploid |  | 0.2034 | 0.6725 |  |
|  | Gain |  |  | 0.9392 |  |
| *YWHAZ* |  |  |  |  |  |
|  |  | Diploid | Gain | Amplification |  |
|  | Shallow Deletion | **0.0021** | **<0.0001** | **<0.0001** |  |
|  | Diploid |  | **<0.0001** | **<0.0001** |  |
|  | Gain |  |  | **<0.0001** |  |

| **Breast Cancer** | | | | | |
| --- | --- | --- | --- | --- | --- |
| **Gene Name** | **CNA Type** | **Number of Cases** | **Mean RNA expression** | **Median RNA expression** | **Standard Deviation** |
| *CCNE2* | Shallow Deletion | 28 | 200.2409964 | 143.995 | 168.1575831 |
|  | Diploid | 360 | 176.3737292 | 133.879 | 214.703652 |
|  | Gain | 504 | 412.1256361 | 297.933 | 408.6577106 |
|  | Amplification | 102 | 517.0755578 | 391.6855 | 446.6727378 |
| *EI3FE* | Shallow Deletion | 27 | 3890.37037 | 3153.4 | 1727.460713 |
|  | Diploid | 354 | 4886.79274 | 4567.395 | 2008.0662 |
|  | Gain | 502 | 7249.936594 | 6459.76 | 3895.633876 |
|  | Amplification | 110 | 9618.729909 | 9083.185 | 3826.01656 |
| *MTDH* | Shallow Deletion | 29 | 2693.94931 | 2640.43 | 628.7879424 |
|  | Diploid | 359 | 3210.547382 | 3191.38 | 853.2903225 |
|  | Gain | 505 | 5431.458356 | 4824.97 | 2501.317747 |
|  | Amplification | 101 | 7251.223267 | 6464.43 | 3361.957177 |
| *MYC* | Shallow Deletion | 25 | 1261.79184 | 1041.24 | 1058.711313 |
|  | Diploid | 340 | 1767.257137 | 1309.455 | 1624.195554 |
|  | Gain | 473 | 2193.311108 | 1745.09 | 1863.139304 |
|  | Amplification | 156 | 3101.409308 | 2317.895 | 2849.032409 |
| *RAD21* | Shallow Deletion | 25 | 4310.7624 | 4082.61 | 1767.969986 |
|  | Diploid | 349 | 5537.876989 | 5160.04 | 2373.51578 |
|  | Gain | 481 | 10365.28102 | 8715.66 | 5927.261345 |
|  | Amplification | 138 | 16311.40804 | 13786.3 | 9799.481356 |
| *TP53INP1* | Shallow Deletion | 28 | 1512.227536 | 1351.23 | 899.4683291 |
|  | Diploid | 360 | 1968.294428 | 1837.165 | 1034.390123 |
|  | Gain | 507 | 2382.528653 | 1934.21 | 2149.760947 |
|  | Amplification | 99 | 3038.233475 | 2405 | 2183.909495 |
| *YWHAZ* | Shallow Deletion | 26 | 12643.49231 | 11975.75 | 4434.917247 |
|  | Diploid | 357 | 15337.30669 | 14301.4 | 4719.271244 |
|  | Gain | 498 | 29067.29279 | 26282.2 | 14456.73131 |
|  | Amplification | 112 | 39314.53482 | 36589.55 | 18258.86377 |
| **Gene Name** | **CNA TYPE** | **CNA TYPE COMPARED (P-Values)** | | | |
| *CCNE2* |  |  |  |  |  |
|  |  | Diploid | Gain | Amplification |  |
|  | Shallow Deletion | 0.6045 | **<0.0001** | **<0.0001** |  |
|  | Diploid |  | **<0.0001** | **<0.0001** |  |
|  | Gain |  |  | **0.0063** |  |
| *EI3FE* |  |  |  |  |  |
|  |  | Diploid | Gain | Amplification |  |
|  | Shallow Deletion | **0.0003** | **<0.0001** | **<0.0001** |  |
|  | Diploid |  | **<0.0001** | **<0.0001** |  |
|  | Gain |  |  | **<0.0001** |  |
| *MTDH* |  |  |  |  |  |
|  |  | Diploid | Gain | Amplification |  |
|  | Shallow Deletion | **0.001** | **<0.0001** | **<0.0001** |  |
|  | Diploid |  | **<0.0001** | **<0.0001** |  |
|  | Gain |  |  | **<0.0001** |  |
| *MYC* |  |  |  |  |  |
|  |  | Diploid | Gain | Amplification |  |
|  | Shallow Deletion | 0.1149 | **0.0007** | **<0.0001** |  |
|  | Diploid |  | **<0.0001** | **<0.0001** |  |
|  | Gain |  |  | **<0.0001** |  |
| *RAD21* |  |  |  |  |  |
|  |  | Diploid | Gain | Amplification |  |
|  | Shallow Deletion | **0.0007** | **<0.0001** | **<0.0001** |  |
|  | Diploid |  | **<0.0001** | **<0.0001** |  |
|  | Gain |  |  | **<0.0001** |  |
| *TP53INP1* |  |  |  |  |  |
|  |  | Diploid | Gain | Amplification |  |
|  | Shallow Deletion | **0.024** | **0.0164** | **0.0004** |  |
|  | Diploid |  | 0.2218 | **0.0001** |  |
|  | Gain |  |  | **0.0021** |  |
| *YWHAZ* |  |  |  |  |  |
|  |  | Diploid | Gain | Amplification |  |
|  | Shallow Deletion | **0.0017** | **<0.0001** | **<0.0001** |  |
|  | Diploid |  | **<0.0001** | **<0.0001** |  |
|  | Gain |  |  | **<0.0001** |  |

**Supplementary Table 3.** Enrichment of specific pathways based on genes of interest.

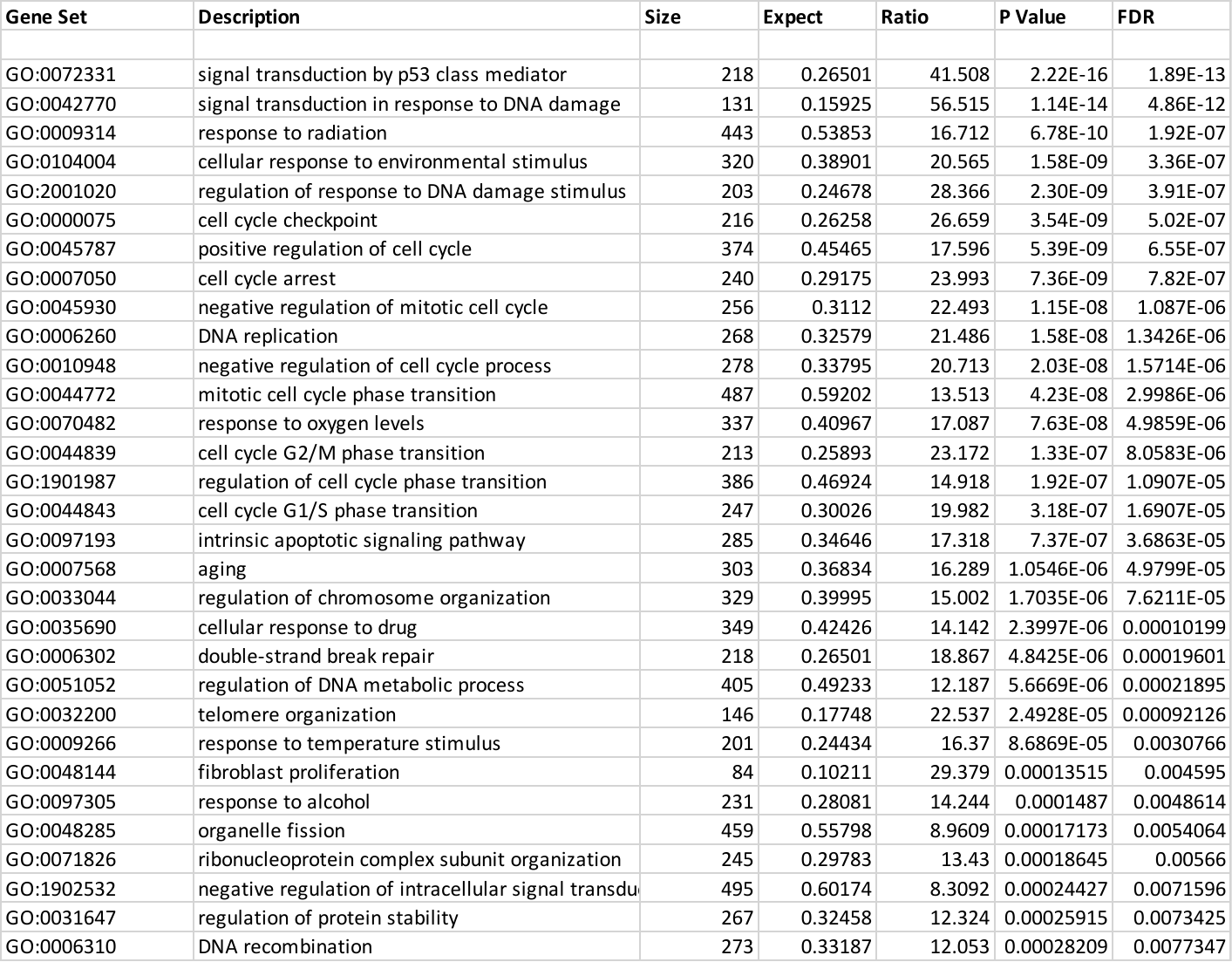

**Supplementary table 4.** Kaplan-Meier curves statistics.

| **Ovarian cancer (OV)** | | | |
| --- | --- | --- | --- |
| **OS** | | | |
| Comparisons | *MYC* and *RRM2B* | *MYC* | *RRM2B* |
| Unaltered | 0.2286 | 0.014 | 0.9055 |
| *MYC* and *RRM2B* |  | 0.0019 | 0.8349 |
| *MYC* |  |  | 0.3653 |
| **DFS** | | | |
| Comparisons | *MYC* and *RRM2B* | *MYC* | *RRM2B* |
| Unaltered | 0.7553 | 0.3125 | 0.8662 |
| *MYC* and *RRM2B* |  | 0.2182 | 0.973 |
| *MYC* |  |  | 0.5174 |
| **Breast Cancer (BRCA)** | | | |
| **OS** | | | |
| Comparisons | *MYC* and *RRM2B* | *MYC* | *RRM2B* |
| Unaltered | 0.1065 | 0.4823 | 0.2966 |
| *MYC* and *RRM2B* |  | 0.8142 | 0.5972 |
| *MYC* |  |  | 0.6729 |
| **DFS** | | | |
| Comparisons | *MYC* and *RRM2B* | *MYC* | *RRM2B* |
| Unaltered | 0.4012 | <0.001 | 0.266 |
| *MYC* and *RRM2B* |  | <0.001 | 0.5684 |
| *MYC* |  |  | 0.0519 |
| **Lung Cancer (LUAD)** | | | |
| **OS** | | | |
| Comparisons | *MYC* and *RRM2B* | *MYC* | *RRM2B* |
| Unaltered | 0.8855 | 0.2717 | 0.835 |
| *MYC* and *RRM2B* |  | 0.3687 | 0.6097 |
| *MYC* |  |  | 0.8166 |
| **DFS** | | | |
| Comparisons | *MYC* and *RRM2B* | *MYC* | *RRM2B* |
| Unaltered | 0.4189 | 0.0734 | 0.0057 |
| *MYC* and *RRM2B* |  | 0.8074 | 0.3372 |
| *MYC* |  |  | 0.2758 |
| **Head and Neck Cancer (HNSC)** | | | |
| **OS** | | | |
| Comparisons | *MYC* and *RRM2B* | *MYC* | *RRM2B* |
| Unaltered | 0.3181 | 0.1726 | 0.4238 |
| *MYC* and *RRM2B* |  | 0.8799 | 0.5392 |
| *MYC* |  |  | 0.5778 |
| **DFS** | | | |
| Comparisons | *MYC* and *RRM2B* | *MYC* | *RRM2B* |
| Unaltered | 0.9766 | 0.2157 | 0.6151 |
| *MYC* and *RRM2B* |  | 0.386 | 0.6399 |
| *MYC* |  |  | 0.5046 |

**Supplementary Table 5.** Tumor Mutation Burden (TMB) profile in OV, BRCA and HNSC.

| **No RRM2B**  **Amplification** | **RRM2B Amplifications** |
| --- | --- |
| Average Mut/MB | Average Mut/MB |
| **OV** |  |
| **48.49586777** | **50.4057971** |
| **BRCA** |  |
| **61.28963795** | **63.04938272** |
| **HNSC** |  |
| **149.1226216** | **147.516129** |

**Supplementary Table 6.** Mutation Signature statistics for BRCA and LUAD.

| **Breast cancer** |  |  |  |  |  |
| --- | --- | --- | --- | --- | --- |
| **Point mutation** | **Trinucleotide** | **One-way ANOVA (*RRM2B* vs others)** | **Two-way ANOVA (*RRM2B*)** | **Two-way ANOVA (*MYC*)** | **Wilcoxon rank-sum** |
| C>A | ACA | 0.132286 | 0.072177 | 0.980320 | 0.631947 |
| C>A | ACC | 0.846501 | 0.996503 | 0.154677 | 0.618010 |
| C>A | ACG | 0.106782 | 0.776010 | 0.817884 | 0.824616 |
| C>A | ACT | 0.558708 | 0.776010 | 0.547911 | 0.303486 |
| C>A | CCA | 0.490546 | 0.559791 | 0.980320 | 0.618010 |
| C>A | CCC | 0.737151 | 0.537633 | 0.982217 | 0.618010 |
| C>A | CCG | 0.713269 | 0.996503 | 0.982217 | 0.561095 |
| C>A | CCT | 0.662572 | 0.713233 | 0.915115 | 0.303486 |
| C>A | GCA | 0.846501 | 0.400952 | 0.154677 | 0.730093 |
| C>A | GCC | 0.799268 | 0.516670 | 0.207840 | 0.303486 |
| C>A | GCG | 0.846501 | 0.996503 | 0.262485 | 0.872521 |
| C>A | GCT | 0.490546 | 0.976837 | 0.873769 | 0.204192 |
| C>A | TCA | 0.821988 | 0.908989 | 0.980038 | 0.637760 |
| C>A | TCC | 0.977458 | 0.996503 | 0.949933 | 0.618010 |
| C>A | TCG | 0.737782 | 0.973564 | 0.980320 | 0.556471 |
| C>A | TCT | 0.940711 | 0.859240 | 0.980320 | 0.952827 |
| C>G | ACA | 0.930462 | 0.833857 | **0.002510** | 0.719065 |
| C>G | ACC | **0.034723** | 0.305316 | 0.255648 | 0.204192 |
| C>G | ACG | 0.308203 | 0.596857 | **0.007232** | 0.618010 |
| C>G | ACT | 0.859373 | 0.976837 | 0.322190 | 0.917894 |
| C>G | CCA | 0.612773 | 0.596857 | 0.817884 | 0.364987 |
| C>G | CCC | **0.034723** | 0.833857 | 0.423385 | 0.303486 |
| C>G | CCG | 0.837624 | 0.996503 | **0.025376** | 0.952827 |
| C>G | CCT | 0.737151 | 0.634226 | **0.002510** | 0.719065 |
| C>G | GCA | 0.675680 | 0.908989 | **0.009984** | 0.618010 |
| C>G | GCC | 0.208639 | 0.196583 | 0.980038 | 0.507435 |
| C>G | GCG | 0.759823 | 0.472805 | 0.510946 | 0.561095 |
| C>G | GCT | 0.633291 | 0.996503 | **0.009984** | 0.368987 |
| C>G | TCA | 0.836912 | 0.908989 | 0.980320 | 0.577262 |
| C>G | TCC | 0.836912 | 0.973564 | 0.982217 | 0.556471 |
| C>G | TCG | 0.846501 | 0.909367 | 0.982217 | 0.863188 |
| C>G | TCT | 0.821988 | 0.850322 | 0.982217 | 0.637760 |
| C>T | ACA | 0.737151 | 0.833857 | 0.980038 | 0.303486 |
| C>T | ACC | 0.675680 | 0.971522 | 0.980320 | 0.470170 |
| C>T | ACG | 0.456929 | 0.537633 | 0.817884 | 0.303486 |
| C>T | ACT | 0.930462 | 0.976837 | 0.980038 | 0.618010 |
| C>T | CCA | 0.936050 | 0.996503 | 0.980320 | 0.637760 |
| C>T | CCC | 0.737151 | 0.976837 | 0.980038 | 0.303486 |
| C>T | CCG | 0.610532 | 0.622146 | 0.817884 | 0.616344 |
| C>T | CCT | 0.882104 | 0.997989 | 0.980038 | 0.824616 |
| C>T | GCA | 0.846501 | 0.908989 | 0.980038 | 0.364987 |
| C>T | GCC | 0.846501 | 0.906358 | 0.980038 | 0.446989 |
| C>T | GCG | 0.502743 | 0.596857 | 0.831368 | 0.556471 |
| C>T | GCT | 0.859373 | 0.921229 | 0.980038 | 0.637760 |
| C>T | TCA | 0.737151 | 0.833857 | 0.980038 | 0.523727 |
| C>T | TCC | 0.713269 | 0.859240 | 0.980038 | 0.594670 |
| C>T | TCG | 0.662572 | 0.833857 | 0.980038 | 0.303486 |
| C>T | TCT | 0.662572 | 0.833857 | 0.980038 | 0.303486 |
| T>A | ATA | **0.000006** | 0.158863 | 0.913911 | 0.618010 |
| T>A | ATC | 0.558708 | 0.586696 | 0.817884 | 0.618010 |
| T>A | ATG | **0.002492** | **0.014568** | 0.364966 | 0.691766 |
| T>A | ATT | **0.000000** | **0.024297** | 0.519866 | 0.413023 |
| T>A | CTA | **0.000045** | 0.158863 | 0.980320 | 0.303486 |
| T>A | CTC | **0.000251** | 0.101769 | 0.980038 | **0.009168** |
| T>A | CTG | **0.008457** | 0.142697 | 0.817884 | 0.656248 |
| T>A | CTT | 0.138434 | 0.559791 | 0.171919 | 0.561095 |
| T>A | GTA | **0.000000** | **0.007032** | 0.577772 | 0.204192 |
| T>A | GTC | 0.432907 | 0.996503 | 0.519866 | 0.719065 |
| T>A | GTG | **0.000381** | 0.577301 | 0.364966 | 0.460507 |
| T>A | GTT | **0.003520** | 0.596857 | 0.817884 | 0.204192 |
| T>A | TTA | **0.000902** | **0.014568** | 0.785995 | 0.335263 |
| T>A | TTC | **0.007767** | 0.569441 | 0.845620 | 0.303486 |
| T>A | TTG | **0.000002** | 0.091287 | 0.980038 | 0.204192 |
| T>A | TTT | **0.000001** | **0.029968** | 0.982217 | 0.303486 |
| T>C | ATA | **0.000003** | **0.014568** | 0.262485 | 0.303486 |
| T>C | ATC | **0.001207** | 0.474474 | 0.980320 | 0.934139 |
| T>C | ATG | **0.000010** | **0.014568** | 0.322190 | 0.364987 |
| T>C | ATT | **0.016114** | 0.586696 | 0.980038 | 0.719065 |
| T>C | CTA | **0.000251** | 0.053394 | 0.364966 | 0.303486 |
| T>C | CTC | **0.000902** | 0.055759 | 0.785995 | 0.637760 |
| T>C | CTG | **0.000576** | 0.053394 | 0.364966 | 0.481995 |
| T>C | CTT | **0.000004** | **0.024297** | 0.720792 | 0.335263 |
| T>C | GTA | **0.000000** | **0.007032** | 0.154677 | 0.364987 |
| T>C | GTC | **0.001933** | **0.049204** | 0.579357 | 0.719065 |
| T>C | GTG | **0.001907** | 0.053394 | 0.364966 | 0.637760 |
| T>C | GTT | **0.000000** | **0.006528** | 0.165370 | 0.108384 |
| T>C | TTA | 0.200897 | 0.453927 | 0.915115 | 0.884800 |
| T>C | TTC | 0.241280 | 0.584808 | 0.980038 | 0.574265 |
| T>C | TTG | 0.182240 | 0.279284 | 0.915115 | 0.561095 |
| T>C | TTT | **0.004188** | 0.067600 | 0.982217 | 0.561095 |
| T>G | ATA | 0.490546 | 0.973564 | 0.692242 | 0.879243 |
| T>G | ATC | 0.066595 | 0.118580 | 0.980320 | 0.204192 |
| T>G | ATG | 0.351968 | 0.537633 | 0.345815 | 0.364987 |
| T>G | ATT | **0.012529** | 0.971522 | 0.817884 | 0.952827 |
| T>G | CTA | 0.280189 | 0.976837 | 0.212997 | 0.303486 |
| T>G | CTC | **0.003116** | 0.250494 | 0.817884 | 0.668654 |
| T>G | CTG | **0.000251** | 0.142697 | 0.980038 | 0.364987 |
| T>G | CTT | **0.003693** | 0.196583 | 0.913911 | 0.364987 |
| T>G | GTA | **0.000725** | 0.081250 | 0.980038 | 0.193376 |
| T>G | GTC | 0.846501 | 0.996503 | 0.262485 | 0.906533 |
| T>G | GTG | **0.002040** | **0.024297** | 0.547911 | 0.303486 |
| T>G | GTT | **0.000156** | **0.006528** | 0.980038 | 0.660001 |
| T>G | TTA | 0.084359 | 0.106510 | 0.982217 | 0.236208 |
| T>G | TTC | 0.573918 | 0.272349 | 0.817884 | 0.556471 |
| T>G | TTG | 0.710637 | 0.842339 | 0.207840 | 0.879409 |
| T>G | TTT | **0.004931** | 0.161062 | 0.929341 | 0.561095 |
| **NSCLC** |  |  |  |  |  |
| **Point mutation2** | **Trinucleotide3** | **One-way ANOVA (RRM2B vs others)4** | **Two-way ANOVA (RRM2B)5** | **Two-way ANOVA (MYC)6** | **Wilcoxon rank-sum7** |
| C>A | ACA | **0.000795** | 0.111467 | 0.692464 | 0.108747 |
| C>A | ACC | **0.000212** | 0.060052 | 0.692464 | 0.059537 |
| C>A | ACG | **0.015558** | 0.082295 | 0.822699 | 0.089879 |
| C>A | ACT | **0.000103** | **0.025644** | 0.753740 | 0.070532 |
| C>A | CCA | **0.000000** | **0.000460** | 0.259552 | 0.059888 |
| C>A | CCC | **0.000000** | **0.003686** | 0.510288 | 0.059537 |
| C>A | CCG | **0.000000** | **0.003456** | 0.259552 | 0.059537 |
| C>A | CCT | **0.000000** | **0.000240** | 0.259552 | 0.059537 |
| C>A | GCA | **0.000621** | 0.054438 | 0.721094 | 0.074688 |
| C>A | GCC | **0.003533** | 0.060052 | 0.822699 | 0.080519 |
| C>A | GCG | **0.001457** | 0.054438 | 0.692464 | 0.059537 |
| C>A | GCT | **0.000598** | 0.082295 | 0.908713 | 0.059537 |
| C>A | TCA | **0.000137** | 0.054438 | 0.692464 | 0.080519 |
| C>A | TCC | **0.000212** | 0.054438 | 0.692464 | 0.059537 |
| C>A | TCG | **0.000310** | **0.035808** | 0.621205 | 0.070532 |
| C>A | TCT | **0.000001** | **0.003456** | 0.510288 | 0.070532 |
| C>G | ACA | **0.028365** | 0.392489 | 0.987396 | 0.150412 |
| C>G | ACC | **0.000465** | 0.134173 | 0.987396 | 0.252147 |
| C>G | ACG | 0.928710 | 0.605241 | 0.848806 | 0.628530 |
| C>G | ACT | **0.034498** | 0.054438 | 0.908713 | 0.074688 |
| C>G | CCA | **0.001051** | 0.185683 | 0.987396 | 0.059537 |
| C>G | CCC | **0.007183** | 0.185683 | 0.987396 | 0.070532 |
| C>G | CCG | **0.001326** | 0.080706 | 0.692464 | 0.070532 |
| C>G | CCT | **0.040347** | 0.134173 | 0.908713 | 0.175647 |
| C>G | GCA | **0.019170** | 0.403219 | 0.908713 | 0.059537 |
| C>G | GCC | **0.001310** | **0.035808** | 0.497736 | 0.087993 |
| C>G | GCG | **0.000062** | 0.054438 | 0.510288 | 0.059537 |
| C>G | GCT | 0.364109 | 0.578460 | 0.908713 | 0.260388 |
| C>G | TCA | 0.253875 | 0.445103 | 0.987396 | 0.138867 |
| C>G | TCC | **0.028365** | 0.204539 | 0.908713 | 0.059537 |
| C>G | TCG | 0.083830 | 0.445103 | 0.990139 | 0.121362 |
| C>G | TCT | 0.429018 | 0.574027 | 0.987396 | 0.150412 |
| C>T | ACA | **0.023421** | 0.273786 | 0.987707 | 0.093232 |
| C>T | ACC | 0.279064 | 0.585507 | 0.987396 | 0.121877 |
| C>T | ACG | 0.240500 | 0.367598 | 0.974327 | 0.089498 |
| C>T | ACT | **0.045257** | 0.711007 | 0.974327 | 0.139044 |
| C>T | CCA | 0.062820 | 0.303764 | 0.908713 | 0.150412 |
| C>T | CCC | 0.083830 | 0.445103 | 0.987396 | 0.166388 |
| C>T | CCG | 0.163448 | 0.445103 | 0.993173 | 0.367509 |
| C>T | CCT | **0.030574** | 0.314535 | 0.987396 | 0.070532 |
| C>T | GCA | 0.292133 | 0.605241 | 0.987396 | 0.226733 |
| C>T | GCC | 0.253824 | 0.369148 | 0.987396 | 0.263822 |
| C>T | GCG | 0.505252 | 0.196601 | 0.987396 | 0.114644 |
| C>T | GCT | 0.071033 | 0.323939 | 0.822699 | 0.289005 |
| C>T | TCA | 0.327887 | 0.586579 | 0.987396 | 0.467022 |
| C>T | TCC | 0.090368 | 0.228788 | 0.987396 | 0.138867 |
| C>T | TCG | 0.833235 | 0.835295 | 0.908713 | 0.882896 |
| C>T | TCT | 0.093298 | 0.271139 | 0.990139 | 0.089498 |
| T>A | ATA | **0.028365** | 0.117898 | 0.987396 | 0.162361 |
| T>A | ATC | 0.152848 | 0.091761 | 0.702321 | 0.688398 |
| T>A | ATG | **0.000586** | 0.082295 | 0.908713 | 0.087993 |
| T>A | ATT | 0.496148 | 0.344784 | 0.987396 | 0.291192 |
| T>A | CTA | **0.012183** | 0.060052 | 0.822699 | 0.074688 |
| T>A | CTC | **0.000212** | **0.025644** | 0.692464 | 0.086529 |
| T>A | CTG | **0.011285** | 0.170428 | 0.692464 | 0.089498 |
| T>A | CTT | **0.008957** | 0.185683 | 0.908713 | 0.138867 |
| T>A | GTA | 0.060145 | 0.228788 | 0.987396 | 0.173248 |
| T>A | GTC | **0.002480** | 0.058853 | 0.822699 | 0.138867 |
| T>A | GTG | **0.007183** | 0.088774 | 0.868845 | 0.089498 |
| T>A | GTT | **0.019328** | 0.228788 | 0.987396 | 0.070532 |
| T>A | TTA | **0.001849** | 0.095001 | 0.895237 | 0.059537 |
| T>A | TTC | **0.007475** | 0.108372 | 0.753740 | 0.070532 |
| T>A | TTG | **0.037218** | 0.266978 | 0.987396 | 0.449633 |
| T>A | TTT | **0.036995** | 0.181837 | 0.987396 | 0.470119 |
| T>C | ATA | **0.036858** | 0.111614 | 0.987396 | 0.184705 |
| T>C | ATC | **0.045181** | **0.025644** | 0.987396 | 0.070532 |
| T>C | ATG | **0.006362** | 0.054438 | 0.987396 | **0.045216** |
| T>C | ATT | 0.192820 | 0.819879 | 0.987396 | 0.121877 |
| T>C | CTA | **0.028365** | 0.076078 | 0.974327 | 0.189785 |
| T>C | CTC | **0.018472** | 0.134173 | 0.908713 | **0.045216** |
| T>C | CTG | 0.084312 | 0.206340 | 0.990139 | 0.108747 |
| T>C | CTT | 0.246343 | 0.228788 | 0.987396 | 0.812004 |
| T>C | GTA | 0.102194 | 0.060052 | 0.987396 | 0.275727 |
| T>C | GTC | 0.453416 | 0.192552 | 0.987396 | 0.783357 |
| T>C | GTG | 0.503065 | 0.159264 | 0.987707 | 0.252147 |
| T>C | GTT | 0.115475 | 0.111614 | 0.987396 | 0.071239 |
| T>C | TTA | 0.320080 | 0.152832 | 0.915802 | 0.150595 |
| T>C | TTC | 0.719414 | 0.192552 | 0.987396 | 0.362539 |
| T>C | TTG | 0.136893 | 0.185683 | 0.987396 | 0.059537 |
| T>C | TTT | 0.307949 | 0.291695 | 0.987396 | 0.182170 |
| T>G | ATA | 0.465491 | 0.882500 | 0.987396 | 0.467022 |
| T>G | ATC | 0.928710 | 0.461149 | 0.908713 | 0.691966 |
| T>G | ATG | **0.026774** | 0.574027 | 0.987396 | 0.628530 |
| T>G | ATT | 0.968682 | 0.862758 | 0.510288 | 0.943552 |
| T>G | CTA | 0.105978 | 0.400010 | 0.908713 | 0.191324 |
| T>G | CTC | 0.172540 | 0.228788 | 0.987396 | 0.111373 |
| T>G | CTG | 0.128043 | 0.206340 | 0.987396 | 0.335674 |
| T>G | CTT | 0.623842 | 0.906204 | 0.997838 | 0.074688 |
| T>G | GTA | **0.000212** | 0.111614 | 0.692464 | 0.059537 |
| T>G | GTC | 0.375213 | 0.323939 | 0.987707 | 0.121362 |
| T>G | GTG | **0.028365** | 0.185683 | 0.974327 | 0.059537 |
| T>G | GTT | 0.783886 | 0.835295 | 0.987396 | 0.520871 |
| T>G | TTA | **0.046279** | 0.060052 | 0.908713 | 0.248212 |
| T>G | TTC | 0.783886 | 0.857651 | 0.987396 | 0.362539 |
| T>G | TTG | 0.071033 | 0.060052 | 0.848806 | 0.384331 |
| T>G | TTT | 0.697048 | 0.835295 | 0.987396 | 0.439276 |
